# A self-organizing single-cell morphology circuit optimizes *Podophrya collini* predatory trap structure

**DOI:** 10.1101/2025.11.12.688106

**Authors:** Zhejing Xu, Lauren E. Mazurkiewicz, Marine Olivetta, Marisol Morrissey, Omaya Dudin, Amy M. Weeks, Scott M. Coyle

## Abstract

Cellular structure self-organizes through an interplay between internal mechanisms and external cues. In predatory ciliates, structures that capture prey obtain the resources needed for their construction, creating feedback between environmental inputs and morphological outputs. Here, we describe a self-organizing single-cell morphology circuit that adaptively optimizes the predatory trap structure of the suctorian *Podophrya collini*. These trap structures ensnare large cellular targets using an array of straw-like tentacles that siphon out prey cytoplasm upon contact with their tips. We find that trap architecture scales anisotropically, favoring tentacle number over length, to construct traps that maximize capture probability for the resources on hand. Drug perturbations, transcriptomics, proteomics, and expansion microscopy define distinct molecular and structural requirements that regulate trap structure maintenance and tentacle biogenesis. We integrate these findings into a mathematical model that explains the adaptive scaling of the trap and that makes predictions we confirm experimentally. More broadly, this circuit’s architecture provides a general-purpose control logic for organizing the number and size of sub-cellular structures applicable to other natural and engineered cellular systems.

## Introduction

Cells possess a remarkable capability to build microscopic structures, dynamically patterning a pool of nanoscale building blocks into diverse target architectures whose forms and functions are critical for biology^1–6^. Such assemblies are organized by the interaction between internal regulation and external cues, allowing structures to be dynamically constructed and disassembled adaptively^7–11^. For example, leading-edge motility organizes a dynamic front and contractile rear to the cell, which orients in response to different environmental stimuli^2,12–15^ . Similarly, neuron morphology is dynamically constructed during outgrowth to tune the number of synapses and the length of the branching neurites^16–19^. However, understanding the specific structure/function relationships at play and the circuitry that directs morphological organization can be challenging in complex multicellular settings.

Single-celled organisms provide intuitive models for studying feedback between cellular structure and the environment^20–25^ . For example, many predatory ciliates build elaborate cellular structures dedicated to prey capture and feeding, whose architectures are readily scored and whose functions are well defined^3,20,26^. These include *Lacrymaria olor*, which searches locally for nearby prey through rapid cycles of extension and contraction of a neck-like proboscis^27–29^; and *Didinium*, which swims rapidly through its environment, wielding a cone-like protrusion on its snout that engulfs *Paramecium* prey upon collision^30–32^. For these cells, the architecture of the predatory structure is generally fixed within the species and scales isotropically with cell size.

In contrast, suctorian ciliates are sit-and-wait predators, capturing prey using trap structures whose morphology and architecture vary across individuals from the same species^33–35^. Among the suctoria, *Podophrya collini* traps large *Tetrahymena* preys using an array of microtubule-based feeding tentacles that radiate out from a spherical body to detect, paralyze, and feed on prey that contact their tips^36–38^ (Fig. 1A, S1A). This creates feedback between the structure of the trap and the prey density of the environment, as the cell’s tentacles are assembled using resources acquired from capture events (Fig. 1B). In routinely fed lab populations, we find that *P. collini* traps display a broad range of tentacle numbers, from as few as one to more than twenty. Tentacle lengths are comparatively constrained, ranging from ∼15-35 μm (Fig. 1E).

**Figure 1.**
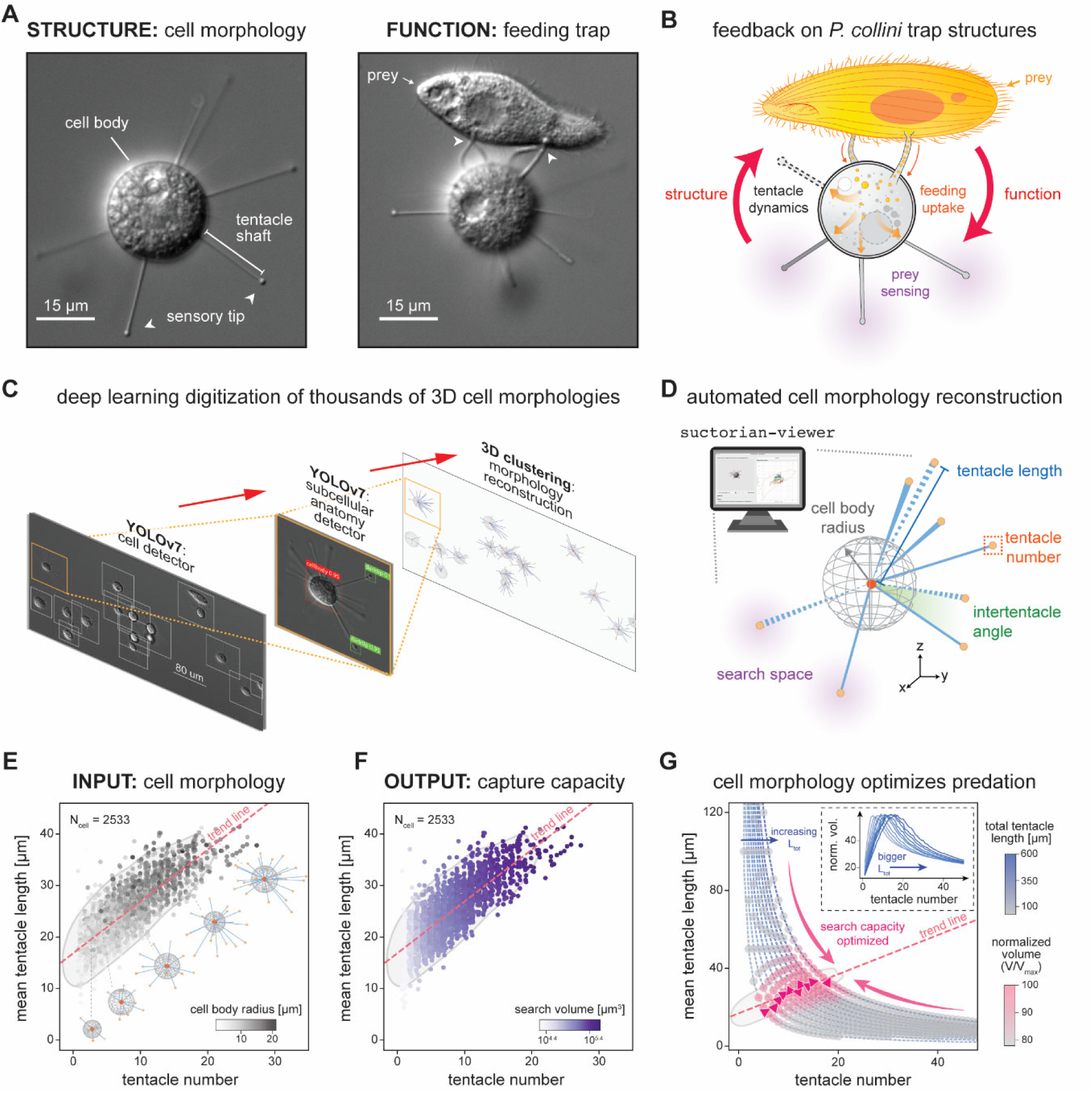
The single-celled suctorian *P. collini* captures prey using tentacle trap structures that scale anisotropically to optimize feeding and predation. (A) Representative bright-field DIC images of a *P. collini* cell during resting and feeding states. In the resting state, the cell extends multiple feeding tentacle traps into the surrounding environment while remaining sessile. Upon collision of prey such as *Tetrahymena pyriformis* (shown) with the tentacle tips (white arrows), *P. collini* rapidly recognizes and captures the prey within milliseconds, with subsequent nutrient flux transported through the tentacles into the cell body. (B) Schematic illustrating the feedback control by which *P. collini*’s structural morphology determines its predatory functions, including feeding uptake and prey recognition. (C) Automated high-throughput cell digitization pipeline for reconstructing *P. collini’s* three-dimensional geometry. DIC Z-stack images were collected, and the maximum-intensity projection was analyzed using a deep-learning-based algorithm (YOLO v7. Detected cells were processed for second-tier YOLO-based identification of cell body and tentacle tips, and their spatial coordinates were extracted. Three-dimensional clustering of these features enables digitization and reconstruction of individual *P. collini* cell morphologies. (D) Representative 3D cell morphology reconstruction corresponding to the cell shown in (A). *P. collini’*s cell geometry was visualized using a custom-built *suctorian-viewer* software. Cell geometric information, including cell body radius, tentacle number and length, and its spatial distribution, can further be extracted. See Fig. S1 for additional detail. (E) Mapping of population-level tentacle configuration in the morphological parameter space (“*morphospace*”) reveals variations in cell body sizes and anisotropic scaling of tentacles. Each cell is plotted by its tentacle number and mean tentacle length, colored by cell body radius (n = 2533 cells). A linear trendline between the tentacle number and mean tentacle length is indicated (pink dashed line). (F) The 3D search space of *P. collini* defines its capture capacity based on individual cell morphology. For each cell, the search volume was estimated as the non-overlapping union of prey-sized (r_search_=17.5 μm) spheres surrounding the tentacle tips. The resulting search capacity was projected onto the morphospace and color-coded on a log_10_ scale. The linear fitting trendline between the tentacle number and mean tentacle length is indicated (pink dashed line). (G) Normalized search volume estimation reveals optimization of tentacle structural allocation. Theoretical search volumes were computed for all possible resource allocations of total length L_tot_ (50-600 μm) between the number and the mean tentacle length, averaged over a range of prey-sized search radii (15-20 μm). Each configuration’s search volume was normalized to the maximum achieved for that L_tot_ and mapped onto the morphospace. Experimentally observed configurations are shown as a 2D kernel density plot (outer contour at 0.05), overlaid with the empirical trend in tentacle scaling. Inset: normalized search volume as a function of tentacle number.

Here, we study *P. collini* trap structure variation quantitatively and systematically using a combination of 3D live-cell morphology digitization and tracking, transcriptomics, proteomics, expansion microscopy, and mathematical modeling. This uncovers an underlying single-cell morphology circuit that acts to optimize *P. collini*’s trap structure for predation. Internally, this circuit sets a specific anisotropic scaling of the cell’s tentacle configuration that distributes resources in a way that maximizes the search volume a trap covers. Externally, the trap’s capture performance feeds back into the cell’s resource pool, steering tentacle configurations towards the best possible structure compatible with the resources on hand. We clarify the circuit’s implementation by disentangling separate molecular and structural requirements at play for tentacle maintenance and biogenesis. This allowed us to develop a mathematical model that both explains the observed adaptive trap scaling behavior and makes predictions about perturbation outcomes that we could confirm experimentally. More generally, the circuit’s architecture provides a universal control logic that can organize the number and size of any cellular sub-structures, translatable to diverse suctorian species as well as other natural and engineered cellular systems.

## Results

### *P. collini* tentacle trap structures scale anisotropically to optimize feeding and predation

The suctorian *P. collini* captures prey using tentacle trap structures that display diverse configurations (tentacle number and length) within the population. To systematically study this variation, we built a deep-learning pipeline that digitizes the 3D morphology of *P. collini* cells. A custom two-tier YOLO^39,40^ detector identifies all *P. collini* in a DIC Z-stack, then uses 3D point clustering to localize sub-cellular structures and extract a complete cell morphology, including the number, length, and spatial organization of the cell’s tentacles (Fig. 1C-1E and Fig. S1B-1D). This allows us to digitize thousands of high-resolution *P. collini* tentacle trap structures from a single experiment without fixation for visualization and analysis in 3D with the accompanying custom-built *suctorian-viewer* software app.

Using this pipeline, we visualized the variation in tentacle trap configuration using a *morphospace*^41^ that plots tentacle number against average tentacle length for each cell (Fig. 1E). Trap structures within the population occupied a specific sub-region of the morphospace, with tentacle number and length appearing to scale in a correlated but anisotropic manner, biased towards tentacle number. Individual tentacle lengths within a trap were generally distributed closely to the trap’s mean tentacle length and uniformly across the cell surface (Fig. S1B-D). A linear fit of the morphospace explains 66% of the population variation (n = 2533 cells; Fig. 1D) and provides a blueprint for how *P. collini*’s trap structure scales: the minimal trap is a single ∼18 μm tentacle and each additional tentacle correlates with a small increase in average tentacle length (∼1.0 μm / tentacle). Relative to this minimum configuration, tentacle lengths can increase by up to 2.3-fold while tentacle number can increase by more than 25-fold.

Because capture occurs upon prey contact with a tentacle tip^37^, we asked how these different tentacle configurations would impact predation. We modeled a trap structure’s predatory capacity as a three-dimensional search volume defined by the union of prey-sized spheres about each tentacle tip^42,43^. In this framework, a cell morphology is an INPUT with an associated OUTPUT search volume. For *Tetrahymena*-sized preys (r_prey_ ∼ 15-20 μm), projecting an experimental morphology’s predatory capacity onto the morphospace revealed that increases in search volume are primarily associated with increases in tentacle number (Fig. 1F). Cells with sparse tentacle configurations benefit the most from additional tentacles, since each tip provides a new, non-overlapping point in space at which capture can occur. However, once tentacles are dense, further gains in search volume also require increases in tentacle length to reduce overlap between tips (Fig. S1E). While larger trap structures cover a greater search volume, they are also more expensive to build, both in terms of their total cost (integrated tentacle length) but also in terms of the cost-efficiency of the strategy (area searched per unit cost) (Fig. S1C and Fig. S1F). Given this tension, how should cells distribute resources between tentacle length and number to maximize the trap’s predatory capacity? We used our search-space model to compare the performance of empirically observed morphologies to other theoretical tentacle configurations. For any cellular resource level *L_tot_*, we can calculate an associated search volume for all possible resource allocations *{(N, L): N×L=L_tot,_}* to identify the optimal trap configuration. Remarkably, the empirically observed morphologies generated near-optimal search volumes across all resource levels tested, with the theoretical best performers aligning closely to the centerline of the *P. collini* morphospace (Fig. 1G). Thus, the anisotropic scaling of *P. collini* tentacle configuration appears to distribute resources and organize the trap structure optimally for capture of *Tetrahymena* prey.

### The *P. collini* morphospace adapts to changing prey density

The anisotropic morphology scaling we identified in routinely fed *P. collini* populations generates tentacle trap configurations that appear to optimize resource allocation for prey capture. As a result, prey density might provide feedback that naturally steers the population towards the most effective predatory strategies achievable under the current environmental conditions. To measure the plasticity of the *P. collini* morphospace, we tracked populational-level changes over a 30-day feed-starve-refeed cycle (Fig. 2A, Fig. S2A-C). Feeding and starvation altered the distribution of cell morphologies within the population, stretching and contracting the accessible regions of the morphospace along the anisotropic scaling line we identified previously (∼1.0 μm per tentacle, R^2^ = 0.669, n = 16,989; Fig. 2B, 2C). Notably, these morphological changes were reversible: refeeding the 28-day starved population distributed cells across the full extent of the morphospace within 24 hours (Fig. 2D).

**Figure 2.**
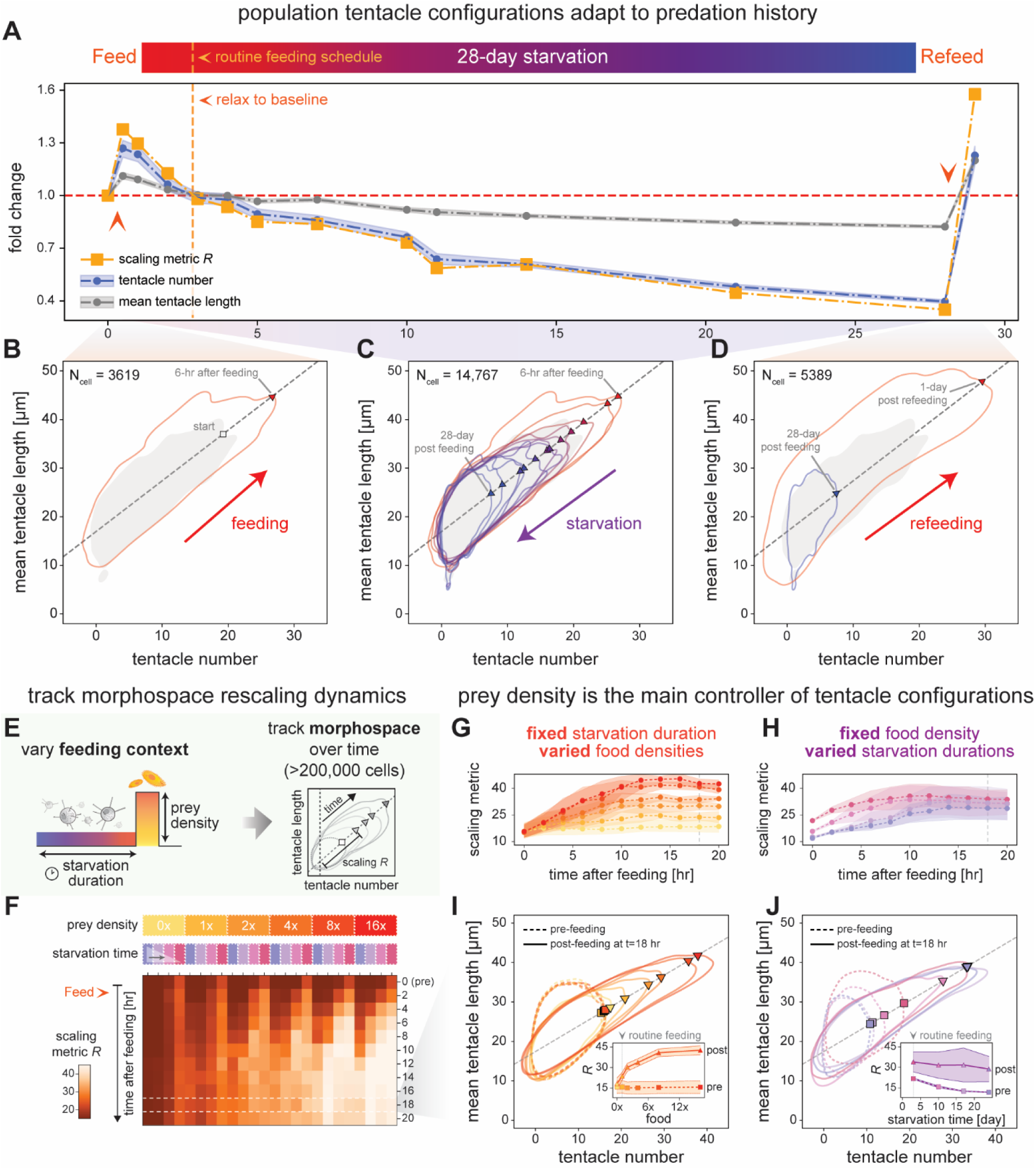
*P. collini* trap structure morphospace adapts to changing prey density. (A) *P. collini* population tentacle configurations change in response to a 30-day feed-starve-refeed cycle. Cells were collected from routinely fed cultures (t_0_) and tracked over time. The fold-change in population-averaged tentacle configurations relative to the baseline feeding condition (t_0_) was plotted over time (n_tot_ = 17,195 cells): tentacle number (blue), mean tentacle length (gray), scaling metric *R* (orange, defined below). Shaded envelopes indicate the population standard deviation. (B-D) Population tentacle configuration distributions shift across the 30-day feed-starve-refeed cycle. Experimentally observed configurations at each time point are shown as a 2D kernel density plot (outer contour at 0.05) overlaid on the baseline morphospace (t_0_, gray). The overall linear fit between the tentacle number and mean tentacle length is indicated (gray dashed line). Morphospace boundary intersections with the scaling line are marked with symbols (square for t_0_ and triangles for subsequent time points). (B) Feeding induces expansion of the morphospace boundary. (C) Prolonged starvation causes gradual boundary contraction. (D) Refeeding restores the morphospace boundary. (E) Experimental design for tracking population-scale rescaling dynamics under varying feeding contexts. *P. collini* populations were harvested after different starvation durations (4, 10, 17, and 24 days) and refed with prey at different densities (0x, 1x, 2x, 4x, 8x, and 16x relative to routine feeding). Populations (n_tot_ = 243,875 cells) were continuously tracked for 20 hours before and after feeding, and sampled every 2 hours. (F) Time-resolved population changes in scaling metric *R* across varying feeding contexts, where population tentacle configurations vary primarily as a function of prey density. The scaling metric *R* quantifies overall population-level scaling response, defined as the distance from the minimal cell morphology (tentacle number = 1) to the intersection of the morphospace boundary with the scaling line. (G, I) Tentacle configuration dynamics under varying food densities with fixed starvation time. Top: Scaling metric dynamics over time (dots: mean, shaded envelopes: standard deviation). Bottom: Morphospace comparison between pre-feeding (dashed line) and 18-hour post-feeding (solid line) populations. Inset: Changes in scaling metric *R* at these two time points across different prey densities. (H, J) Tentacle configuration dynamics under varying starvation durations with fixed food density. Top: Scaling metric dynamics over time (dots: mean, shaded envelopes: standard deviation). Bottom: Morphospace comparison between pre-feeding (dashed line) and 18-hour post-feeding (solid line) populations. Inset: Changes in scaling metric *R* at these two time points across different starvation durations.

To compactly quantify and summarize changes in population morphology over time, we define a scaling metric *R,* as the distance from the minimal cell morphology to the morphospace boundary’s intersection with the scaling line. Plotting *R* across the feed-starve-refeed cycle recapitulates the observed morphological transitions, with the morphospace expanding in scale upon feeding and reducing in scale upon starvation (Fig. 2A). Thus, prey availability appears to dynamically control what regions of the morphospace are accessible, while the anisotropic nature of this scaling ensures that the resulting trap configurations always remain optimal for predation. To further disentangle the contributions of starvation history and prey density to the rescaling of the morphospace, we starved *P. collini* for different lengths of time (3, 10, 17, 24 days) and tracked the morphospace dynamics upon refeeding with a range of prey densities (every 2 hours over 20 hours, n = 243,876) (Fig. 2E, 2F, Fig. S2D-H). We found that the morphospace scales directly in proportion to prey density (Fig. 2G, 2I). Prey density was the primary controller, as populations with radically different starvation histories ultimately converged to similar scales when fed the same prey density (Fig. 2I, 2J), albeit with different recovery kinetics (Fig. 2G, 2H). Thus, the *P. collini* morphospace dynamically rescales to adapt to changing prey conditions. Intriguingly, the scaling parameter *R* can be viewed as a population estimate of prey availability: the more expanded the morphospace, the higher the prey density, and *vice versa*.

### Cells actively remodel their tentacle configuration to tune structure to resource availability

The *P. collini* population morphospace dynamically scales in response to changes in prey density, resulting in a distribution of tentacle trap structures that are optimal for prey capture. Such remodeling could arise from selective processes acting on the population; alternatively, individual cells might be able to adapt their tentacle configurations directly. To distinguish these possibilities, we developed an agar-based microchamber culturing setup that enables longitudinal observation of individual *P. collini* cells over weeks (Fig. 3A)^44,45^. Confining *Tetrahymena* and *P. collini* in the same microchamber results in heterogeneous feeding outcomes, such that only some cells feed successfully. Using our analysis pipeline, we digitized and tracked single-cell morphology trajectories in response to prey capture and prolonged starvation.

**Figure 3.**
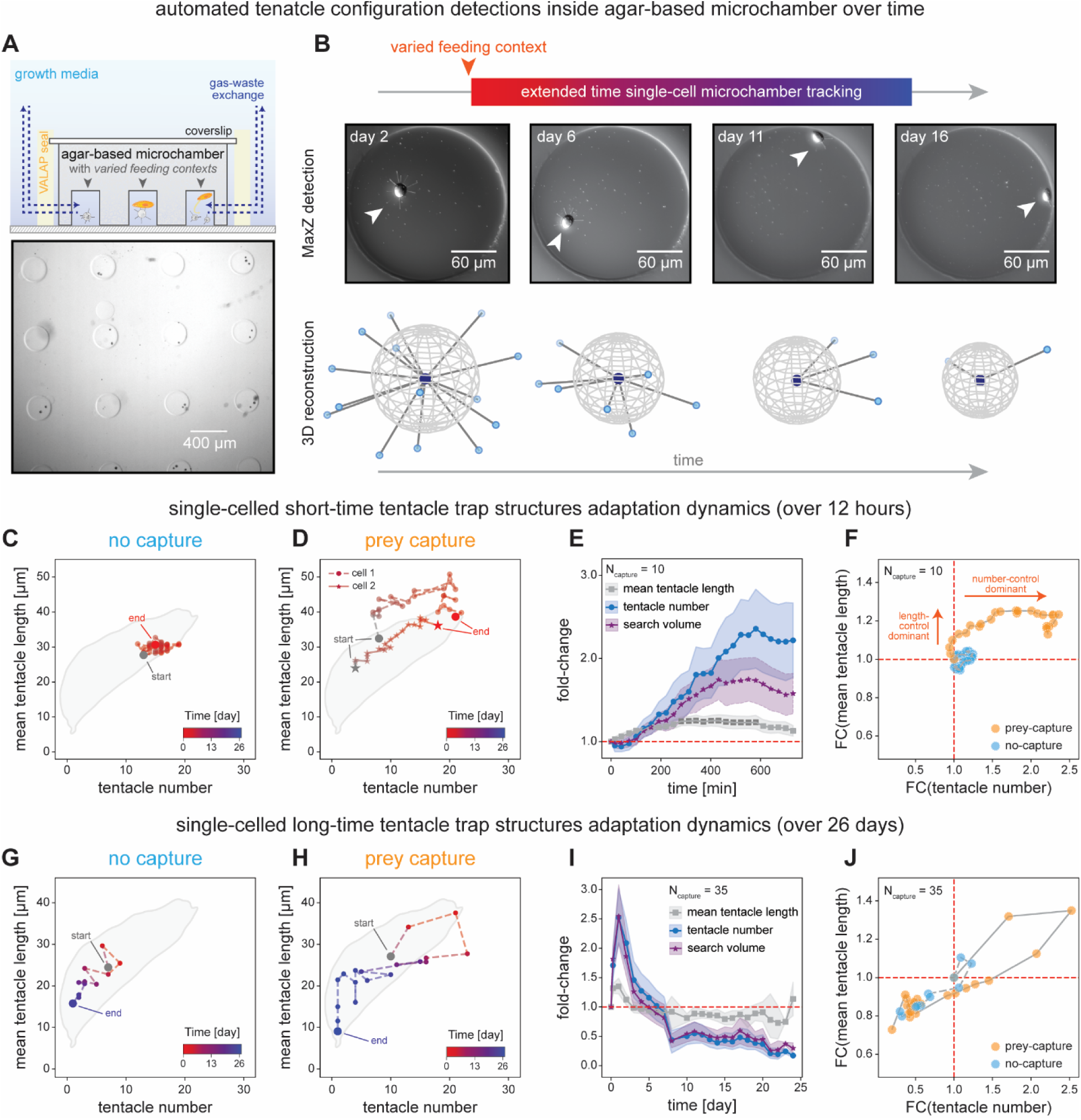
Individual *P. collini* cells actively remodel their tentacle configuration to tune trap structure to resource availability. (A) A modified agar-based arrayed microchamber system enables long-term single-cell live-cell tracking. Molding 3-5% Nobel agar with a PDMS stamp produces 250 μm diameter microchambers with a 250 μm chamber depth. Individual *P. collini* and *T. pyriformis* are mixed and confined within single chambers, which are sealed with VALAP for attachment and surrounded by growth medium. Bottom: bright-field image of *P. collini* cells confined in arrayed microchambers. (B) Automated tracking of individual *P. collini* tentacle configuration trajectories over time. Cells in different feeding contexts were imaged in 3D to monitor temporal changes in cell morphology. Shown are representative maximum intensity projection images (top) and corresponding 3D reconstructions (bottom). (C-F) Short-term (12-hour) adaptation dynamics of single-cell trajectories over time. (C) Representative trajectory of a single-cell tentacle configuration without prey capture. Initial and final configurations are indicated. (D) Representative trajectories of two cells, each with one successful prey capture in its own microchamber. The initial time point corresponds to prey release (end of feeding). (E) Averaged fold-change trajectories of single-cell tentacle parameters for captured cells (n_capture_ = 10 cells). Symbols represent mean values; shaded regions denote standard deviation. Gray: mean tentacle length; blue: tentacle number; purple: search volume. (F) Scatterplot showing relative fold-changes in tentacle number and mean tentacle length among single cells with or without capture (n_nocapture_ = 9 cells). (G-J) Long-term (26-day) adaptation dynamics of single-cell trajectories over time. (G) Representative trajectory of a cell without prey capture. (H) Representative trajectory of a cell with one successful prey capture. Cells shown in (G) and (H) were confined in the same microchamber. Initial and final configurations are indicated. (I) Averaged fold-change trajectories of tentacle parameters for captured cells (n_capture_ = 35 cells). (F) Scatterplot of relative fold-changes in tentacle number and mean tentacle length among single cells with or without capture (n_nocapture_ = 11 cells).

In the absence of prey, cells showed minimal change in the trap configuration overnight, with tentacle number and length fluctuating by only 7.3% and 3.1%, respectively (n = 9; Fig. S3A). In contrast, cells that successfully captured prey underwent dramatic and coordinated morphological remodeling. On average, end-state tentacle number increased 2.4-fold and tentacle length increased 1.2-fold after capture, expanding the trap’s search volume nearly two-fold (n = 10; Fig. 3E, 3F). Importantly, these changes are a direct result of prey capture, not soluble cues: in chambers containing multiple *P. collini*, only the cells that successfully captured prey expanded their morphology (Fig. 3G, 3H). Thus, successful predation provides new resources that trigger the anisotropic expansion of tentacle configuration at a single cell level.

To understand how cells remodel their trap structures, we separately tracked tentacle length and number dynamics following prey capture. Tentacle elongation was rapid and immediate, reaching a new steady state within ∼100 minutes (t_0.5_ ∼ 55 minutes; Fig. 3E). The emergence of new tentacles, however, typically lagged: while some cells initiated new tentacle formation shortly after feeding, most began later, with some taking more than 100 minutes for the first new tentacle to appear (Fig. 3D). After this initial delay, tentacle number increased consistently before stabilizing around 9 hours post-feeding (t_0.5_ ∼ 5.4 hours; Fig. 3E). These dynamics reveal a clear separation of timescales between tentacle length control and biogenesis, hinting at different regulatory mechanisms at play.

We continued to track the longer-term morphological trajectories of individual cells over weeks of starvation. Cells that captured prey and initially expanded their tentacle configurations returned to their baseline morphology within ∼ 4 days (Fig. 3H). These cells continued pruning tentacles to as few as one or two over the next 26 days (n = 34; Fig. 3I, 3J). Cells that did not capture prey followed a similar morphological trajectory but without the initial expansion associated with prey capture (n = 11; Fig. 3G, Fig. S3B). In both cases, starvation trajectories followed the scaling trend of the population-level morphospace downwards towards increasingly minimal morphologies and ultimately death.

These experiments establish that individual *P. collini* cells can actively remodel their tentacle trap structure, expanding it upon prey capture and dismantling it during starvation. These morphological trajectories actively steer cells towards tentacle configurations with resource allocations that optimize search volume. Thus, a self-organizing single-cell morphology circuit must somehow encode the scaling, dynamics, and functional adaptation of the tentacle trap.

### Tentacle maintenance and biogenesis are regulated by different cellular mechanisms

We next sought clarity over the implementation of the adaptive trap-scaling morphology circuit, which depends on a cell’s ability to regulate both the length and number of its tentacles. Because increases in length and number appeared to operate at different timescales, we suspected that distinct molecular processes might govern tentacle maintenance and biogenesis. To explore this possibility, we developed a high-throughput assay for studying synchronous tentacle biogenesis. Using mild pressure homogenization to sever a cell’s tentacles, we could trigger synchronous *de novo* biogenesis across the population with a half-time of ∼ 4 hours (Fig. 4F, S4A, S4B, S4D-K). This “de-tentacle” assay allows us to separately probe the requirements for tentacle biogenesis and maintenance through integrated biochemical, transcriptomic, and proteomic analyses.

**Figure 4.**
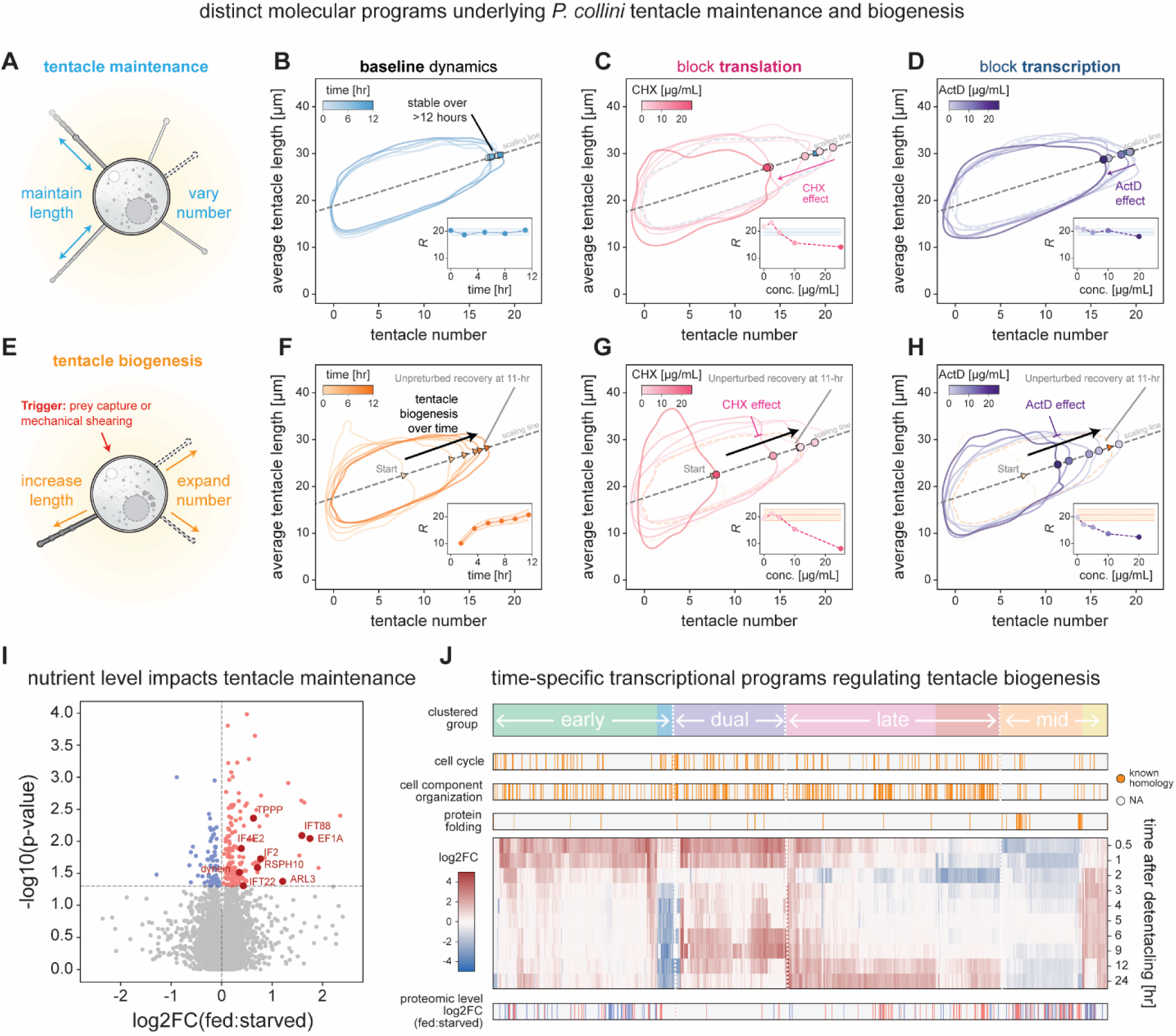
Distinct cellular programs regulate *P. collini* tentacle maintenance and biogenesis. (A-D) Molecular programs underlying tentacle maintenance. (A) Schematic illustrating dynamic controls of tentacle length and number for maintaining the tentacle trap structures. (B) Tentacle configuration changes in routinely fed *P. collini* populations (baseline, n_base,tot_ = 22,818 cells) were tracked over 12 hours and mapped onto the morphospace. Inset: scaling metric *R* over time, overlaid with the population mean ± standard deviation as the baseline reference. (C) Population morphospace response of the routinely fed *P. collini* (n_CHX,11-h_ = 2730 cells) incubated with a gradient of cycloheximide (CHX) concentrations to block translation for 11 hours. Inset: scaling metric R across CHX concentrations, overlaid with the baseline mean ± standard deviation. (D) Population morphospace response of the routinely fed *P. collini* (n_ActD,11-h_ = 2275 cells) incubated with a gradient of actinomycin D (ActD) concentrations to block transcription for 11 hours. Inset: scaling metric *R* across ActD concentrations, overlaid with the baseline mean ± standard deviation. (E-H) Molecular programs underlying tentacle biogenesis. (E) Schematic illustrating tentacle growth in length and number during tentacle biogenesis, triggered by prey capture or mechanical shearing. (F) Tentacle configuration changes during synchronized de novo tentacle biogenesis following detentacling (tentacle-severing assay; see Fig S4A). Detentacled *P. collini* populations (n_detentacled,tot_ = 12,090 cells) were tracked for their tentacle regeneration over 12 hours and mapped onto the morphospace. Inset: scaling metric *R* over time (shaded area: standard deviation). (G) Morphospace response of recovering populations (n_deCHX,11.5-h_ = 3578 cells) incubated with a CHX concentration gradient for 11.5 hours. Inset: scaling metric *R* across CHX concentrations, overlaid with the mean ± standard deviation of DMSO-treated recovering population at 11.5 hours (negative control). (H) Morphospace response of recovering populations (n_deActD,11.5-h_ = 2857 cells) incubated with an ActD concentration gradient for 11.5 hours. Inset: scaling metric *R* over different ActD, overlaid with the mean ± standard deviation of DMSO-treated recovering population at 11.5 hours (negative control). (I) Label-free mass spectrometry quantification (LFQ) of well-fed and starved *P. collini* populations. Proteins of interest and their UniProt identifications are indicated. (J) Differential transcriptomic analysis of synchronized *de novo* tentacle biogenesis. Transcripts showing significant level changes (|log2FC|>1.5, adjusted p-value<0.05) were clustered into four major groups based on cytological observations (see Fig. S4H-K). Functional annotations and corresponding quantitative LFQ comparisons are illustrated.

We first examined the requirements of transcription and translation for maintenance of a tentacle configuration using the chemical inhibitors actinomycin D^46^ and cycloheximide^47^, respectively. *P. collini* populations treated with DMSO (negative control) showed no significant changes in morphospace structure over time (Fig. 4B). Similarly, actinomycin D had minimal effects even at the highest dose tested (Fig. 4D). However, cycloheximide treatment caused a dose-dependent reduction in morphospace scale (Fig. 4C). This suggests that a cell’s existing tentacle configuration depends on active protein synthesis, linking its maintenance to translational capacity—and by extension, resource availability^48–50^.

To further explore how *P. collini* nutritional state impacts protein synthesis and tentacle maintenance, we compared the proteomes of well-fed and starved *P. collini* cultures using label-free mass spectrometry^51^. Well-fed cells exhibited strong upregulation of host-cell translation factors^52^ (EF1, IF1, IF4, etc.), cytoskeleton-related and structural proteins^53,54^ (dynein, TPPP, etc.), as well as many novel proteins of unknown function (Fig. 4I, Table S1). A more robust and active translational machinery can thus support the ongoing synthesis of structural components needed to maintain larger tentacle configurations, helping stabilize a cell’s position in the morphospace.

We next examined the roles of transcription and translation in tentacle biogenesis by tracking the morphological recovery of de-tentacled cells across different perturbations. DMSO-treated populations fully restored their morphospace with a half-time of approximately four hours (Fig. 4F). As expected, blocking translation with cycloheximide inhibited morphospace recovery in a dose-dependent manner, implicating protein synthesis in both tentacle maintenance and formation (Fig. 4G). Critically, actinomycin D also blocked tentacle regeneration in a dose-dependent manner (Fig. 4H), suggesting a role for new transcription in tentacle biogenesis.

To further define how transcription changes during tentacle biogenesis, we constructed a *de novo* transcriptome for *P. collini* and used RNA sequencing to perform a differential expression analysis across multiple time points during a 24-hour regeneration period. In parallel, we performed tubulin immunostaining to visualize and track the timing of tentacle nucleation and biogenesis. By clustering the transcriptional responses and projecting them onto these cytological phases, we identified four major gene clusters with distinct temporal profiles corresponding to sequential stages of tentacle biogenesis (Fig. 4J, Fig. S4H-S4K, Table S2).

Following tentacle shearing, residual tentacle roots were disassembled into free tubulin within ∼ 30 minutes (Fig. S4C, S4I). Cluster 1 genes, upregulated at these early time points, were enriched for factors involved in filament destabilization and protein degradation (Fig. 4J, Table S2). Around two hours, new tentacles began nucleating within the cytosol as pre-tentacle structures (Fig. S4J, S4K). Clusters 2 and 3, corresponding to this phase, were enriched for genes related to cellular organization and localization, including centrin-like EF-hand cytoskeletal proteins, cilia-related and microtubule-associated proteins (dynein-nexin, kinesins, IFTs, and RSPs), as well as many genes without known homology (Fig. 4J, Table S2). Finally, Cluster 4 contained primarily unannotated genes, many of which were also seen as upregulated in the proteomic profiles of fed populations (Fig. 4J). Thus, distinct waves of transcripts appear to align with the appearance and maturation of new tentacle structures.

### Expansion microscopy reveals tentacle substructures essential for function and assembly

Our sequencing and proteomics data suggest that *P. collini’s* microtubule-based tentacles require additional components and structural features for function, adding to the complexity of tentacle biogenesis. Among genes upregulated during biogenesis, the presence of centrin-like EF-hand proteins suggests potential sensory, contractile, or myoneme-like elements within the tentacle^3,28,55–58^. To explore this possibility, we used immunostaining and confocal microscopy to simultaneously visualize tubulin and centrin distributions in the cell (Fig. 5A). The microtubule components of a tentacle are seen to run all along its length and deep into the cell interior. Interestingly, this structure is decorated with two distinct patterns of centrin localization: a bulb-like structure at the tentacle tip; and a collar-like structure where the microtubules pass through the cell cortex.

**Figure 5.**
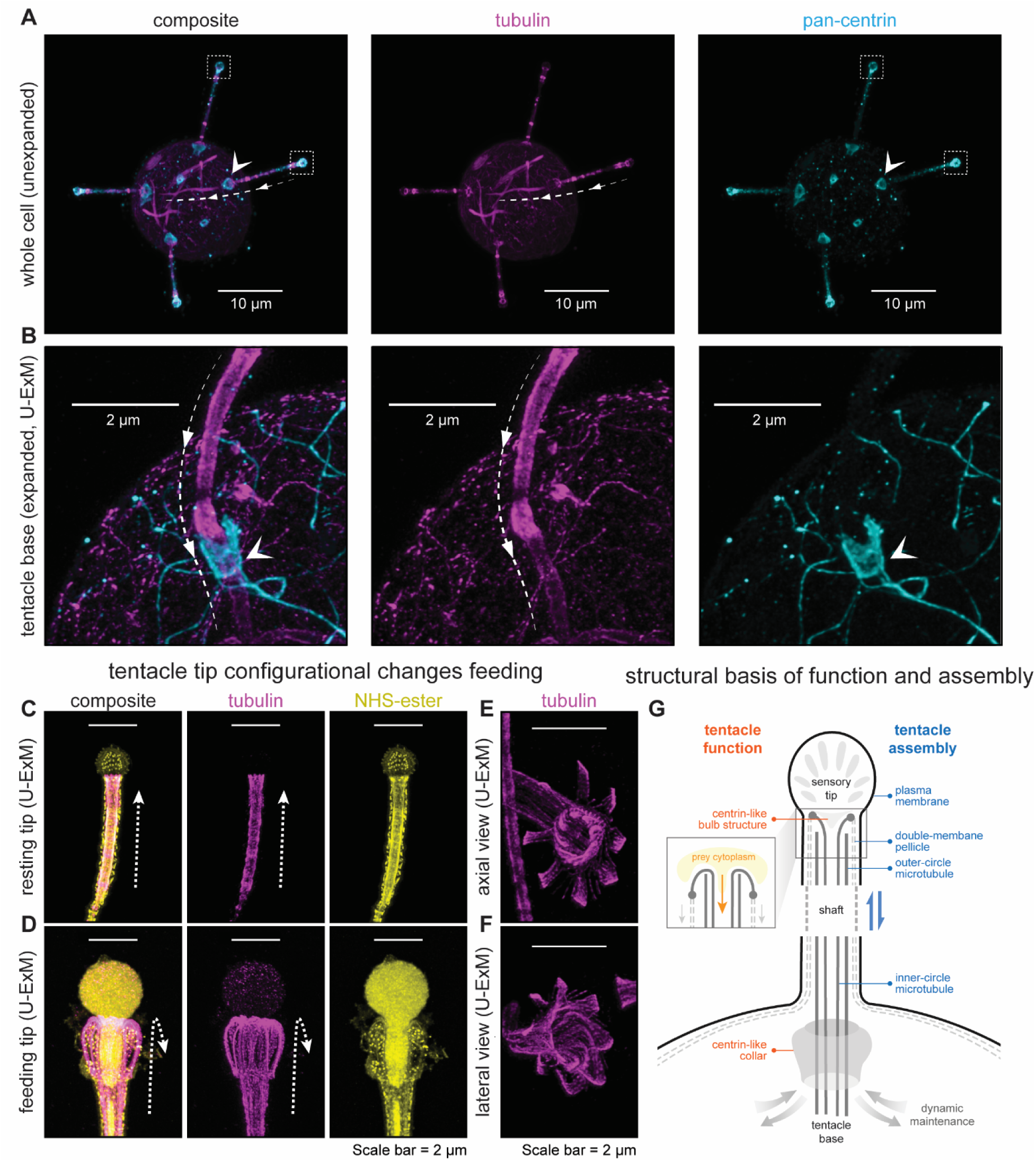
Ultrastructure Expansion Microscopy (U-ExM) reveals *P. collini* tentacle substructures essential for their assembly and function. (A) Representative confocal images of an unexpanded *P. collini* cell immunostained with anti-tubulin (magenta) and anti-centrin (cyan). The dashed line outlines the tentacle microtubule shaft, the arrow indicates the centrin-like collar, and boxes highlight the bulb-like centrin-rich tip. (B) Representative U-ExM images of an expanded *P. collini* cell stained as in (A). The dashed line outlines the tentacle microtubule shaft, and the arrow marks the centrin-like collar in an enlarged view. (C) Representative U-ExM images of a resting *P. collini* tentacle stained with NHS-ester (yellow) and anti-tubulin (magenta). Microtubules appear relaxed along the natural tentacle axis (dashed line). (D) Representative U-ExM images of a feeding *P. collini* tentacle stained as in (C). Microtubules exhibit pronounced bending near the tentacle tip (dashed line). (E, F) Representative U-ExM images showing microtubule (magenta) bending at feeding tentacle tips in (E) axial view and (F) lateral view. (G) Schematics summarizing key substructural features essential for *P. collini* tentacle functions and assembly. Tentacles are composed of an organized microtubule framework enclosed by a pellicular membrane system, with centrin-rich collar and bulb-like tip structures indicating specialized molecular organization. Inset: proposed model of microtubule bending during feeding.

These specialized sub-structures within the tentacle might contribute to prey recognition or feeding activity. To visualize their architecture at higher resolution, we performed Ultrastructure Expansion Microscopy (U-ExM)^3,59–63^, a method increasingly adapted for microbial eukaryotes. Consistent with older EM studies^36,37^, the microtubule scaffolding consists of an inner ring of microtubule ribbons and an outer ring of microtubules (Fig. 5G). In expansion, the centrin collar structure can now be seen to form a hoop that encircles these microtubules, anchoring the tentacle to the cortex while still allowing microtubules to extend further into the cell interior (Fig. 5B). At the tentacle tip, we observed two distinct structural configurations. In the resting state, the microtubule ribbons attached to the cell cortex mark the end of the tentacle shaft, upon which the centrin-rich and NHS-reactive tip structure sits (Fig. 5C). In the active feeding state, the tip structure appears contracted, with the microtubule ribbons splayed open and pulled sharply back (Fig. 5D-5F, Fig. S4L), forming an open feeding gullet to allow for prey consumption.

This ultrastructural analysis provides additional clarity on the structural constraints at play for tentacle maintenance and biogenesis (Fig. 5G). Specialized tip and collar sub-structures with distinct molecular compositions appear to be an essential feature of every tentacle, making their assembly a prerequisite for new tentacle formation. In contrast, the microtubule scaffolding along the tentacle shaft itself appears likely adjustable, such that rapid changes in tentacle length could likely be achieved through microtubule growth or shrinkage.

### Modeling the single-cell morphology circuit that optimizes *P. collini* predatory strategy

Based on these observations, we developed a mathematical model of a morphological circuit that captures the essential features of the resource allocation and feedback regulation that optimizes *P. collini*’s tentacle trap structure. In this model, the cell’s total resource level *R*_tot_is partitioned among *N* tentacles of lengths {*L*_1_, *L*_2_,…, *L*_N_} and a shared free resource pool *R*. These resources correspond to the key building blocks (tubulin, MAPs, etc.) that make up the general tentacle structure and length. Inspired by models of ciliary length control^64–68^, a tentacle’s growth is positively regulated by *R* and negatively regulated by its length; while tentacles shrink at a constant fixed rate. For *N* competing tentacles, mass action defines a steady-state allocation of *R*_tot_ into *N* tentacles of length *L*_ss_, and a free resource pool *R*_ss_ (Fig. 6A; detailed in Methods).

**Figure 6:**
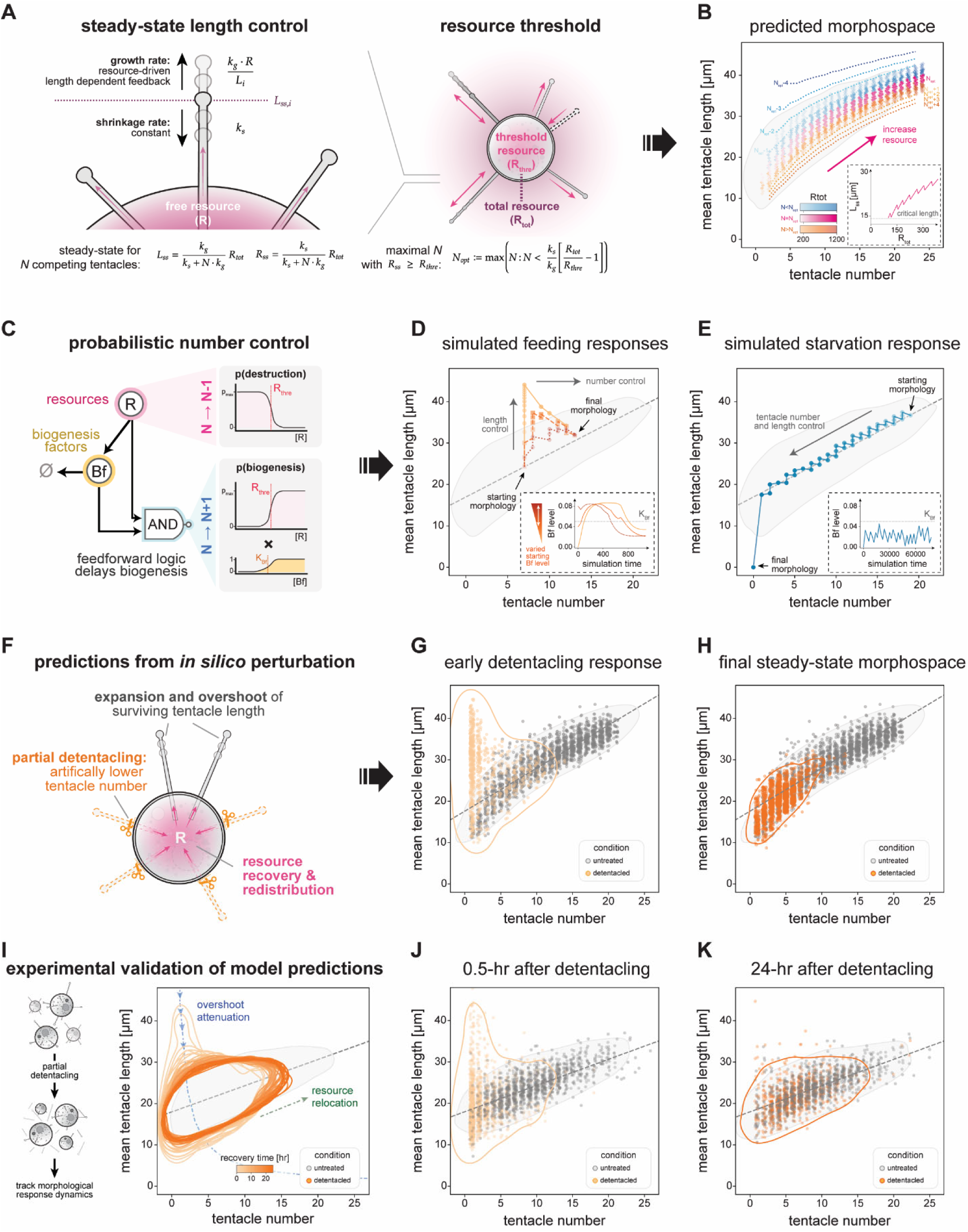
Modeling the single-cell morphology circuit that optimizes *P. collini* trap structure and predatory strategy. (A) Schematics of the resource-controlled steady-state length control module. Left: Individual tentacle length is regulated by deterministic ordinary differential equations describing growth and shrinkage kinetics, yielding a steady-state length *L_ss_* at a given total resource level *R_tot_* and tentacle number *N*. Right: A threshold resource level *R_thre_*, related to *R_tot_*, determines the optimal tentacle number *N_opt_*. (B) Predicted tentacle configurations at increasing *R_tot_*levels based on a logarithmic *R_thre_-R_tot_* relationship. Experimentally observed routinely fed *P. collini* populations (baseline, n_baseline_ = 2,579 cells) define the morphospace boundary (gray, 2D kernel density=0.05). Predicted steady-state tentacle configurations spanned (*N_opt_*±4, *Lss*) are mapped onto the morphospace. Configurations with *N_opt_*are shown in pink; *N_opt_*-n (n=1,2,3,4) in blue; and *N_opt_*+n (n=1,2,3,4) in orange. Inset: Predicted mean tentacle length at *N_opt_* as a function of *R_tot_*. (C) Schematics of the probabilistic number regulation module. Tentacle number dynamics are modeled as stochastic jumps: tentacle destruction depends on resource depletion, and new tentacle formation follows a feedforward process determined by resource levels and biogenesis factors *Bf*. (D) Simulated single-cell feeding responses. Feeding is modeled as a step increase in *R_tot_*. Three cells with identical initial morphologies and resource changes but varying initial *Bf* levels were simulated. Trajectories are mapped onto the baseline morphospace (gray). Inset: *Bf* dynamics over time for three simulated cells (colored by initial *Bf*). (E) Simulated single-cell starvation response. A cell under constant resource drainage was simulated. Its trajectory is mapped onto the baseline morphospace (gray). Inset: *Bf* dynamics over time. (F) Schematics of *in silico* partial detentacling perturbation. The model simulates a sudden partial loss of tentacles and predicts subsequent steady-state adjustments in number and length. (G-H) Simulated population responses to partial detentacling. Predicted steady state of untreated (gray dots, n_sims,un_ = 2000 cells) and partially detentacled populations n_sims,de_ = 2000 cells, nude in G, orange in H) are mapped onto the empirical baseline morphospace (gray). (I) Experimental validation of partial detentacling. Left: schematic of partial detentacling asay. Right: experimental observed population morphospace boundary changes over time (orange, partial detentacled populations N_partialde, total_=58,539 cells; gray, untreated populations N_untreated, total_=58,374 cells). (J) Experimentally observed population tentacle configurations 0.5 hours after partial detentacling. Untreated cells (gray) and partially detentacled (nude) are overlaid on the averaged morphospace of untreated morphospace (gray). (K) Experimentally observed population tentacle configurations 24 hours after partial detentacling. Untreated cells (gray) and partially detentacled (orange) are overlaid on the averaged morphospace of untreated morphospace (gray).

The steady-state tentacle length *L*_ss_ depends on both the total resources *R*_tot_ as well as the number of tentacles in the system, growing longer when *N* is small and shorter when *N* is large. Because short tentacles (<15 μm) were rarely observed, to gain intuition we considered the consequences of a threshold resource quantity *R*_thre_required for tentacle formation (Fig. 6A). Defining *N*_opt_ as the largest number of tentacles satisfying *R*_ss_ > *R*_thre_ , we visualized how (*N*_opt_, *L*_ss_) scales as a function of increasing resources *R*_tot_. Tentacles initially grow smoothly; abruptly decrease when a new tentacle forms; then continue growing as the cycle repeats (Fig. 6B). While a fixed *R*_thre_ only scales tentacle number, a resource-dependent *R*_thre_ reproduces both length and number scaling by providing negative feedback on new tentacle formation (Fig. 6B, S5A-S5D). Although the distribution of ( *N*_opt_, *L*_ss_ ) is much narrower than experimentally observed (Fig. 6B), real populations likely contain cells in transition that have not yet reached their end-state configuration. Indeed, configurations spanned by(*N*_opt_ ± 4, *L*_ss_) effectively captured both morphospace scaling and spread (Fig. 6B), pointing to the need for the morphological circuit to explicitly encode dynamic transitions in number.

To this end, we extended our deterministic length-control model to formally include probabilistic jumps in tentacle number^69^ (Fig. 6C). Tentacle loss occurs stochastically when the free resource level *R* drops below *R*_thre_. To recapitulate delays in biogenesis and its dependency on specific transcriptional products, we modeled it as a feed-forward process^70^: with probabilities determined jointly by *R* and a biogenesis factor *Bf*, whose expression itself is regulated by *R* (Fig. 6C). Numerical simulations for a given *R*_tot_ generate dynamic morphological trajectories that converge to an end-state configuration approaching (*N*_opt_, *L*_ss_), with stochastic fluctuations of ± 1 tentacle (Fig. S6E). Thus, the full dynamic model actively steers a cell’s morphology towards the optimal tentacle configuration for a given resource level.

We used this model to simulate morphological trajectories in response to prey capture, which we modeled as a step increase in total resource level *R*_tot_. As expected, tentacle length and number dynamically adjust to a new steady-state configuration (Fig. 6D). Importantly, a cell’s specific trajectory depends on the initial levels of the biogenesis factor *Bf*: high *Bf* trajectories closely followed the scaling trend, while cells with little or no *Bf* experienced transient overshoots in length before tentacle biogenesis ultimately drove convergence to the same end-state configuration (Fig. 6D). Thus, differences in resting *Bf* levels can potentially explain the diversity of trajectories observed experimentally.

Starvation was simulated as a gradual decline in *R*_tot_, which causes the cell’s tentacles to shrink in length collectively (Fig. 6E). However, stochastic tentacle destruction events transiently replenish free resources, allowing the remaining tentacles to temporarily restore their length and establish a local optimal configuration. Eventually, the system collapses to a state with no tentacles, mirroring the progressive morphological regression observed during prolonged starvation (Fig. 3G, 3H).

Our proposed morphological circuit suggests testable hypotheses about how cells would respond to synthetic perturbations to their trap structures. For example, if the number of tentacles were abruptly reduced, the remaining tentacles should transiently elongate before new tentacles are synthesized (Fig. 6G, 6H). To test, we reduced the applied pressure in our detentacling assay to induce partial tentacle loss within the population. Because residual tentacle roots are recycled upon severing (Fig. S4), this leads to a sudden increase in a cell’s free resource levels. Following perturbation, the population now occupied regions of the morphospace rarely accessed under normal conditions (low *N*, high *L*) but which were predicted to be populated by our model (Fig. 6J). Over time, cells reorganized their tentacle configurations and redistributed back into the expected morphospace boundaries, consistent with the underlying control logic of the morphological circuit (Fig. 6K).

Taken together, a simple resource-based feedback model that incorporates probabilistic tentacle biogenesis and destruction defines a morphological circuit that allows *P. collini* to self-organize trap-structure adaptively to changing resource levels, naturally optimizing predatory strategy for the local environment.

## Discussion

By systematically analyzing the morphological variation and dynamics of *P. collini* trap structures, we identified a self-organizing single-cell morphology circuit that optimizes tentacle configuration for prey capture. This circuit scales the trap anisotropically, biased towards changes in tentacle number over length in a manner suited for efficient capture of *Tetrahymena*-sized prey. During cycles of prey capture and starvation, the circuit steers the trap towards tentacle configurations that maximize capture probability for the available resources. As a result, these single-cell trajectories collectively reorganize the population around the most optimal trap structure that can be effectively maintained by the global prey density of the environment.

Transcriptomic, proteomic, and structural inquiries clarified the implementation of the circuit. Existing tentacles within a trap structure required active translation to be maintained, allowing them to rapidly adjust length in response to resource availability and protein synthesis rates. In contrast, the creation of new tentacle structures appeared to be structurally complex and dependent on resource-sensitive transcriptional programs. This includes the assembly of centrin-containing tip and collar structures that support rearrangement of the tentacle’s microtubule structure during prey sensing and feeding. While additional investigations will be needed to determine the more detailed molecular mechanisms at play, these distinct requirements buffer a cell’s trap configuration against sudden changes in tentacle number until some critical amount of time, resources, and biogenesis factors allow tentacle nucleation to occur.

We formalized these observations in a mathematical model in which deterministic growth of tentacles competing for free resources is interrupted by stochastic jumps in tentacle number occurring when resources are above or below a threshold level *R*_thre_ (Fig. 7A). The deterministic component allows existing tentacles to rapidly adjust length to a steady state resource allocation. When this allocation causes free resources to drop below *R*_thre_, destruction is likely, freeing up resources to support the growth of remaining tentacles. When free resources exceed *R*_thre_ , biogenesis can occur but is potentially delayed by the additional requirement for biogenesis factors whose accumulation is resource dependent. Consistent with experiments, this causes the steady-state trap configuration to be specified by resource levels; but its dynamic transitions to be influenced by predatory history through the resting levels of its biogenesis factors.

**Figure 7.**
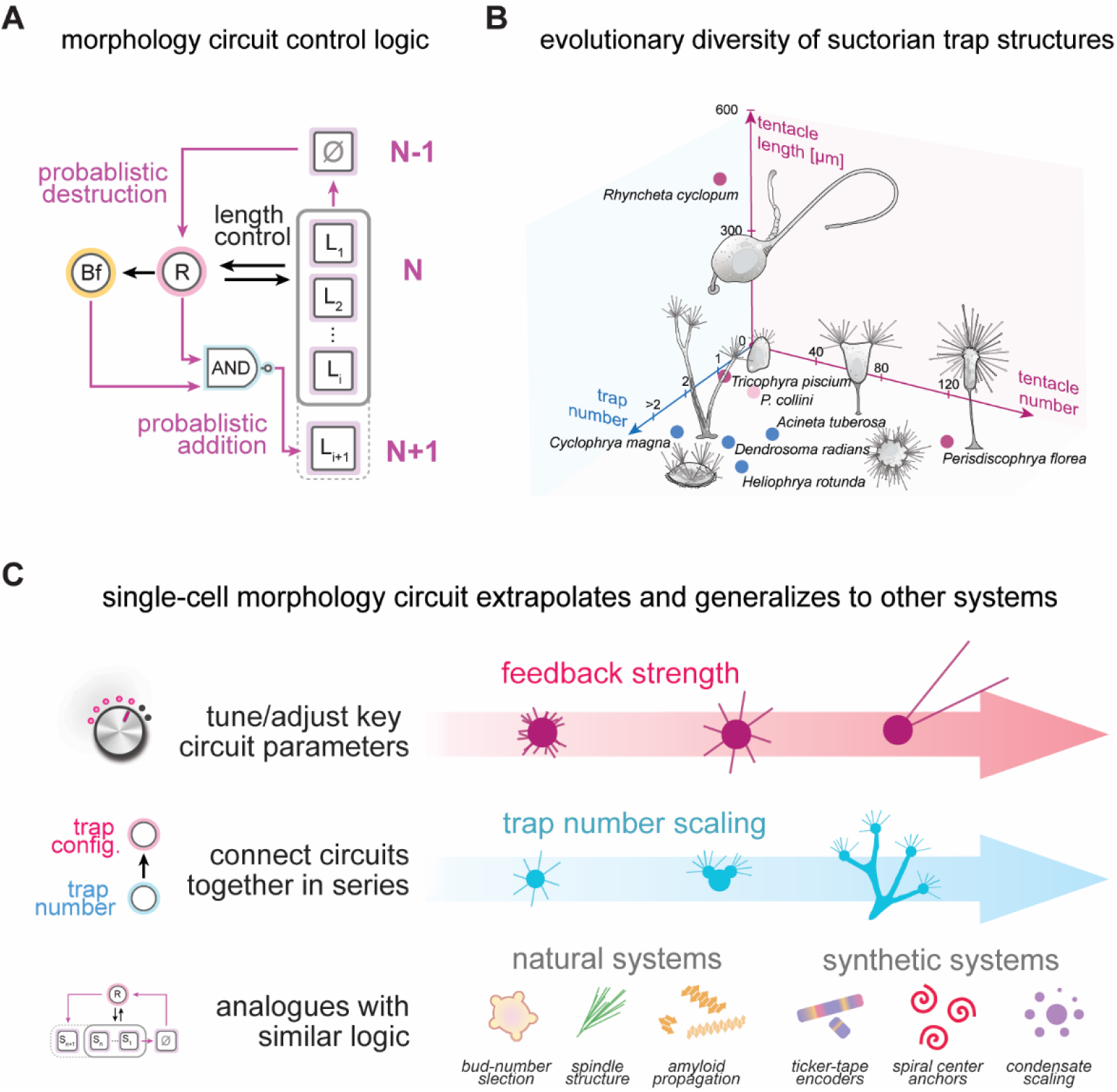
*P. collini* single-cell morphology circuit control logic generalizes to other systems. (A) *P. collini* morphological control circuit. A self-organizing single-cell feedback system integrates deterministic control of tentacle length with stochastic regulation of tentacle number, enabling adaptive optimization of *P. collini* tentacle trap structures. (B) Diverse suctorian trap architectures. Conceptual schematics illustrating how alternative trap configurations with distinct functional trade-offs can emerge from parameter variations or evolutionary extensions of the core circuit. (C) Generalization of the morphological circuit principle. The *P. collini* morphological circuits can be parameter-tuned or connected in series to generate alternative trap strategies. Similar resource-allocation logic, balancing discrete structural numbers and continuous structural sizes, underlies adaptive organization across diverse natural and synthetic systems.

The circuit’s core scaling mechanism arises from biasing stochastic jumps in tentacle number to occur near a free-resource threshold *R*_thre_ . Recapitulating *P. collini*’s specific trap scaling behavior (∼1 μm/tentacle) required the addition of feedback in the model, increasing *R*_thre_as trap structures scale (Fig. S5). As a result, this circuit architecture can encode different scaling behaviors by simply adjusting the feedback strength. Indeed, many trap architectures seen in other suctorian species from different niches can be modeled as different parameterizations of the circuit (Fig. 7B)^35,71–74^ . For example, *Rhyncheta cyclopum* species live as epibionts on crustaceans and typically produce a small number of extremely long tentacles; this configuration is needed to allow their traps to extend far enough away from their hosts to interact effectively with the environment^72^ . Likewise, *Tricophyra* species that colonize the interior of fish gills typically produce a large number of extremely short tentacles; this configuration works for this niche because circulating flows within the gills reduce the effective search burden^35^ .

At the same time, larger suctorian species often build multiple trap structures that are localized to different parts of a larger, more complex cell geometry^71,75^ . While individual traps can be described by our model, extensions are needed to describe how the distribution of multiple trap structures is regulated (Fig. 7B). Interestingly, our proposed morphology circuit can be naturally abstracted to play this higher-level organizing role, scaling trap number and size analogously to how tentacle configurations are scaled within an individual trap. By connecting morphological circuits in series, a cascading resource allocation can in principle be instantiated to couple resource levels to multi-tiered morphological organization in such systems (Fig. 7C).

Indeed, because the function of our proposed morphological circuit is fundamentally to self-organize the distribution of some resource into a discrete number of growth-limited structures, it can potentially be applied to many structure-scaling systems in cell biology (Fig. 7C). For example, multi-budding yeast species map the total amount of Cdc42 in the cell into a discrete number of bud-sites to specify the number of daughter cells they will produce^76^; and condensate-forming proteins in a cellular environment partition into droplet structures whose size and numbers are globally controlled by protein concentration^77–79^. Likewise, the circuit’s logic might be deployable for engineering applications and synthetic structure design. For example, it could be used to tune the number and write-capacity of protein ticker-tape filaments used for recording single-cell transcriptional histories^80,81^; or act to scale the number of spiral cores in programmable reaction diffusion systems or excitable media that serve as sub-cellular anchor points for organizing activities within the cell^82–84^.

By studying how *P. collini* organizes its trap structure morphology, we not only uncovered a mechanism for optimizing predatory strategy in a fascinating single-celled organism but also design principles for adaptive organization of the number and size of subcellular structures more generally. Morphological circuits like this one^2,76,85–91^ can thus be viewed as universal building blocks for organizing cell structure, analogous to how network motifs^92^ and circuit topologies^93,94^ provide conceptual building blocks for organizing gene expression and signaling dynamics.

## Acknowledgments

We thank members of the Coyle Lab, Bill Bement, Jason Cantor, Melissa Harrison, and Phil Newmark for helpful discussions and critical reading of the manuscript.

## Funding

S.M.C. acknowledges support from NSF award 2313723, a David and Lucille Packard Fellowship for Science and Engineering, and USDA/Hatch award WIS03093. A.M.W. acknowledges support from startup funds from the University of Wisconsin-Madison Department of Biochemistry and a David and Lucille Packard Fellowship for Science and Engineering. L.E.M. was supported through the UW-Madison Biotechnology Training Program under grant number NIH 5 T32 GM135066, William H. Peterson Fellowship (University of Wisconsin-Madison Department of Biochemistry), and Dr. Steven Babcock Agricultural Chemistry Graduate Fellowship (University of Wisconsin-Madison Department of Biochemistry). M.O. and O.D. were supported by an SNSF Starting Grant (TMSGI3_218007) and core funding from Department of Biochemistry, University of Geneva. This project is also funded in part by the Gordon and Betty Moore Foundation (GBMF13113) to O.D.

## Data availability

Proteomic and transcriptomic data have been deposited in their appropriate databases. Processed datasets are available on Github and can be viewed using the accompanying suctorian-viewer app. Raw microscopy images and other data types supporting the findings of this study are available from the authors on request.

## Conflict of interest

The authors declare that they have no conflicts of interest with the contents of this article.

## Methods

### Cell line: Podophrya collini and Tetrahymena pyriformis

*Podophrya collini* cell cultures were grown in 30 mL of Volvic mineral water in vented 100 mm x 20 mm cell culture dishes (Greiner 664160). Each culture was fed 0.5 mL of washed *Tetrahymena pyriformis* culture twice per week. *T. pyriformis* cultures were cultured in 10 mL PPY medium (20 g/L protease peptone and 2.5 g/L yeast extract) in 14 mL snap-cap round-bottom test tubes (Falcon) at room temperature in the dark. Details of culture maintenance are described below.

### Detailed culture methods

*P. collini* cultures were obtained as a gift from the Scottish Marine Institute and maintained long-term in the laboratory for experimental use. Culture maintenance protocols were initially developed by Kitching^38^ and modified to increase culture density. Cultures were grown in 30 mL of Volvic mineral water in vented 100 mm x 20 mm cell culture dishes at room temperature in the dark. Each culture was fed 0.5 mL of washed *Tetrahymena* culture twice per week. Over time, bacterial biofilm formation and detritus accumulation occurred naturally as a result of feeding. To maintain culture health and stable population growth, *P. collini* cultures were passed through a 10 μm cell strainer (PluriStrainer) and resuspended in fresh Volvic water every three months.

*T. pyriformis* cultures were also obtained from the Scottish Marine Institute and expanded in bulk as the food source for *P. collini*. Cells were grown in PPY medium (20 g/L protease peptone and 2.5 g/L yeast extract) in 14-mL snap-cap round-bottom test tubes (Falcon) and maintained at room temperature in the dark. Cultures were subcultured weekly to sustain a high-density food source. To initiate a new subculture, 1 mL of culture from the previous week was inoculated into 10 mL of fresh PPY medium.

To prepare *Tetrahymena* for feeding, 2 mL of a two-week-old subculture was washed three times in 13 mL of fresh Volvic mineral water. Cells were pelleted at 1000x RCF for 2 minutes between washes. The final pellet was resuspended in 8 mL of fresh Volvic water, providing sufficient food for four 30 mL *P. collini* cultures for 3-4 days. After feeding, 1-2 mL of *P. collini* culture was removed from each dish to maintain the culture volume and collected for experiments.

### Live cell brightfield microscopy

To concentrate *P. collini* cells and remove detritus, cultures were passed through a 10 μm cell strainer (PluriStrainer) and resuspended in fresh Volvic mineral water. *T. pyriformis* cells were concentrated by centrifugation at 1000x RCF for 2 minutes and washed three times with fresh Volvic mineral water. Both cell types were resuspended in fresh Volvic water for imaging. For imaging, cells were plated either in 96-well glass-bottom plates (Cellvis P96-1.5H-N) for population-level analyses or in 35 mm glass-bottom culture dishes (MatTek P35G-1.5-20-C) for microchamber-based single-cell tracking. Bright-field images were acquired on a Nikon Eclipse Ti2 microscope equipped with a Prime 95B Scientific CMOS (sCMOS) camera using a 10x DIC objective. Given the robustness of our digitization pipelines, cells were also imaged under the same microscope setup with a Kinetix sCMOS camera and a 20x DIC objective. For overnight time-lapse imaging, cells were maintained in a Tokai Hit stage-top incubator set to 25 °C to minimize medium evaporation and volume changes. All brightfield DIC images were collected as Z-stacks at 1 frame per second (FPS).

### Automated 3D cell morphology reconstructions

An automated three-dimensional cell morphology analysis pipeline was developed by integrating a deep-learning-based object-detection model, YOLOv7 (You Only Look Once) v7, with custom Python scripts to detect and reconstruct *P. collini* cell morphology using non-invasive bright-field imaging. The YOLO training dataset is available on the Roboflow project. All YOLO training and detections were performed on GPUs using Google Colab and the Center for High-Throughput Computing (CHTC). Given a collection of bright-field Z-stack image series, a maximum-intensity Z-projection was used to identify and isolate individual cells within each field of view, after which YOLOv7 was applied to crop the corresponding Z-stacks for each detected cell. YOLOv7 was then used to detect the positions of the cell body and individual tentacle tips across the Z-planes, which were converted into XYZ spatial coordinates to represent the tentacle and body geometry. Based on the morphological architecture of *P. collini*, these spatial coordinates were mapped to reconstruct a simplified 3D representation of each cell. This digitization pipeline enables high-throughput, non-invasive quantification of live cell *P. collini* cell morphology and is robust across microscopes with comparable optical configurations.

### Union search volume estimation of *P. collini* trap structure

The three-dimensional search volume of a cell was defined as the non-overlapping union of spheres centered at each tentacle tip, each with the same search radius *r*_search_, representing the cell’s predatory reach. For a cell with *N* tentacles of length {*L*_1_, *L*_2_,…, *L*_i=N_} and a cell body radius *r*_cellbody_, we used Monte Carlo simulations to estimate the search volume by randomly sampling points in the 3D space. Points were uniformly sampled within a bounding box of points large enough to encompass all search spheres. A point was counted as part of the search volume if its distance to any tentacle tip was less than *r*_search_ , and it was located outside the cell body sphere. For empirically observed cells, we used the YOLO-based 3D cell geometry reconstructions to determine cell geometry and estimate the corresponding search volume. For simulated cells, since *P. collini* exhibited nearly uniform tentacle distribution and length, we modeled cells with *N* equally long tentacles (L̄) and assigned cell body radii based on a linear fit from the observed data (*r*_cellbody_∼*f*(*L*_tot_)). The *r*_search_ used in the figures was 17.5 µm, empirically determined based on the size range of *Tetrahymena* preys (15-20 µm).

### Population feed-starve-feed experiment

*P. collini* cells were harvested 3-4 days after their last feeding to establish the baseline population for experiments. The growth medium was refreshed before the experiments by passing cultures through a 10 μm cell strainer (PluriStrainer) and resuspending cells in fresh Volvic mineral water in a new Petri dish (Falcon 351029). After the cells settled, aliquots were plated in a 96-well glass-bottom plate (Cellvis P96-1.5H-N) and imaged as the pre-feeding time point. For the initial feeding, 0.5 mL of washed *Tetrahymena* was added to the master cell culture dish. The cells were imaged at multiple time points following the initial feeding: 6 hours, 1 day, 2 days, 3 days, 4 days, 5 days, 7 days, 10 days, 11 days, 14 days, 21 days, and 28 days. To test whether the *P. collini* morphospace can recover after prolonged starvation, 0.5 mL of washed *Tetrahymena* was reintroduced to the master cell culture dish 28 days post-feeding. The following day, cells were imaged again as part of the refed population. The same experimental workflow was repeated twice, yielding a total of three biological replicates. *P. collini* cell geometry within the populations over time was analyzed using the automated cell morphology reconstruction pipeline.

### Population varying feeding context experiment

*P. collini* cells were starved for four different durations in separate Petri dishes (Falcon 351029). Before the experiment, cultures were cleaned using 10 μm cell strainers (PluriStrainer) and resuspended in fresh Volvic mineral water to ensure consistent cell density and medium conditions. *Tetrahymena* cultures were washed and resuspended in fresh Volvic water = for the experiments. To evaluate the factors contributing to the population-level response in the feed-starve-feed experiment, *P. collini* and *Tetrahymena* cells were combined under varying starvation times (four levels: 3, 10, 17, and 24 days after the last feeding) and prey densities (six levels: 0x, 1x, 2x, 4x, 8x, and 16x, relative to routine feeding). *P. collini* cells were seeded in a 96-well glass-bottom plate (Cellvis P96-1.5H-N) at the same volumetric cell density used for standard culture maintenance. After the cells settled, initial imaging was performed as the pre-feeding time point.

Washed *Tetrahymena* cultures were then added to each volume corresponding to the designated prey densities and starvation times. The plates were imaged every two hours for 20 hours post-feeding. To minimize bacterial growth in the wells, the cells were transferred daily into fresh 96-well plates before imaging. Each feeding context was tested in triplicate. Changes in *P. collini* cell geometry across time and feeding contexts were quantified using the pipeline.

### Individual cell tracking experiment

We adapted an agar-based microchamber setup to track the morphological dynamics of *P. collini* cells over extended periods^44,45^. To confine *P. collini* while maintaining nutrient and fluid exchange, we used a PDMS stamp that generated 250 μm diameter microchambers with a 250 μm chamber depth (Research Micro Stamps). The micropatterned stamp was placed on a single glass slide positioned on a flat surface, and 3-5% liquefied Noble agar was poured onto it. A second slide, oriented perpendicular to the agar and weighted for even pressure, was used to mold the chamber array. The stamp-agar-slide assembly was incubated at room temperature for at least 5 minutes to allow the agar to solidify. After removing the agar mold from the stamp, we transferred the agar pad onto a clean square coverslip (Fisher Scientific 12-541A 18×18-1.5) and trimmed it to fit a 35 mm glass-bottom culture dish (MatTek P35G-1.5-20-C). Each agar pad was loaded with ∼50 μL of cell suspension, which settled for 5-10 minutes to ensure cells occupied the microchambers. Before sealing, the glass-bottom culture dishes were treated with oxygen plasma to enhance adhesion. The agar pad and dishes were then sealed with VALAP (1:1:1 mixture of Vaseline, lanolin, and paraffin). Finally, 3 mL of fresh Volvic mineral water supplemented with 1% penicillin-streptomycin was added to prevent bacterial growth during long-term imaging. To enable prey capture events, *P. collini* and *Tetrahymena* cells were washed in fresh Volvic water, then mixed and loaded onto the agar pad. Individual cells were imaged and tracked over time using the digitization pipeline.

### Tentacle severing and regeneration assay

Baseline *P. collini* cells were washed using 10 μm cell strainers (PluriStrainer) and resuspended in fresh Volvic mineral water. To achieve high-throughput severing, cells were passed through an EmulsiFlex-C3 homogenizer at mild pressure (1000-3000 psi). Detentacled cells were collected by straining the flow-through fraction through 5 μm cell strainers (PluriStrainer). Detached tentacles can be harvested by centrifuging at 6000 x RCF, 4 °C. This approach enables population-level tentacle severing while synchronizing tentacle regrowth over several hours. Tentacle regeneration dynamics were monitored in 96-well glass-bottom plates (Cellvis P96-1.5H-N), and tentacle configuration recovery was quantified either as a detailed 3D tentacle reconstruction or as a binary evaluation (presence/absence of tentacles) using our YOLO pipeline.

### Chemical perturbation experiments

Perturbation experiments were carried out by incubating the cells in fresh Volvic water that contained the indicated concentrations of the perturbant. Two drugs were used to target global cellular processes: actinomycin D (ActD) for transcriptional inhibition and cycloheximide (CHX) for translational inhibition. Concentration gradients were adapted from prior studies: 0, 2.5, 5, 10, and 25 μg/mL for CHX; 0, 2, 5, 10, and 20 μg/mL for ActD. To assess the effects of the drug on tentacle maintenance and tentacle biogenesis, treatments were applied immediately after cell plating. Tentacle configurations were imaged over time in 96-well glass-bottom plates (Cellvis P96-1.5H-N) and analyzed either as a detailed 3D representation of the tentacle configuration or as a binary evaluation (presence/absence of tentacles) using our digitization pipeline.

### Total RNA extraction

Baseline bulk *P. collini* RNA samples were extracted using the Invitrogen PureLink RNA microkit (Invitrogen 12183-016). Cells were strain-cleaned and pelleted by centrifugation at 2000 x RCF for 5 minutes at 4 °C. Cells were lysed with PureLink L3 lysis buffer with dithiothreitol (DTT) and carrier RNA, then homogenized according to the manufacturer’s instructions (Invitrogen 12183–026). RNA extraction was performed following the standard PureLink RNA microkit protocol. The quality of the extracted RNA samples was assessed using NanoDrop (ThermoFisher Scientific) and Agilent RNA chips on a Bioanalyzer (Agilent Technologies). Cells were collected both before and after feeding to construct the bulk *P. collini* transcriptome. Five samples with RNA integrity number (RIN) values above 8.0 were carried for further analysis.

For differential transcriptional profiling during tentacle regeneration, RNA was extracted using the Qiagen RNeasy Micro Kit (Qiagen 74004) due to discontinuity in the PureLink RNA extraction kit between 2020 and 2021. Detentacled cells were strain-cleaned and pelleted by centrifugation at 2000 x RCF for 5 minutes at 4 °C. Cells were lysed in Qiagen RLT buffer containing DTT (without carrier RNA) and homogenized using the Qiagen QIAshredder (Qiagen 79654). Non-detentacled cells from the same experiment were processed in parallel as negative controls. RNA extraction was performed according to the Qiagen RNeasy extraction protocol, and the workflow was repeated to generate three biological replicates. RNA quality was assessed using the NanoDrop (Thermo Fisher Scientific) and the Agilent Bioanalyzer (Agilent Technologies), and samples with RIN values over 8.0 were retained for downstream analysis.

### RNAseq library preparation and sequencing

For the baseline bulk *P. collini* transcriptome, RNA-Seq libraries were prepared using 300 ng of total RNA per sample. Libraries were prepared following the NEBNext Single Cell/Low Input cDNA Synthesis & Amplification kit (New England Biolabs E6421L). cDNA was purified with ProNex beads (Promega NG2001) and amplified with 15 PCR cycles. SMRTbell library preparation of the pooled cDNA fractions was performed using the PacBio SMRTbell Express Template Prep Kit 2.0, following the PacBio Iso-Seq protocol (PN 101-763-900 version 01 (June 2019)). The quality and concentration of the amplified cDNA and final library were assessed using Qubit high-sensitivity (HS) DNA assay (Thermo Fisher Scientific) and Agilent HS DNA chips on a Bioanalyzer (Agilent Technologies). Five libraries were sequenced using three SMRT cells on the PacBio Sequel II platform (PacBio).

For the *P. collini* tentacle regeneration time-course, RNA-Seq libraries were prepared using 200 ng of total input RNA per sample. cDNA libraries were generated using Illumina TruSeq Stranded mRNA (Illumina 20020594) and indexed with IDT for Illumina TruSeq RNA UD Indexes (Illumina 20021454), following the TruSeq Stranded mRNA Reference guide. Library quality and concentrations were analyzed using the Agilent TapeStation (Agilent Technologies). Libraries were sequenced on the NovaSeq6000 platform (Illumina) with a 2×150 bp paired-end reads, reaching an average read depth of 50 million read pairs per detentacled time point library and 200 million read pairs per negative control library.

### *De novo* transcriptome assembly and annotations

PacBio Iso-Seq raw data were processed using IsoSeq3 v3.3.0 on a local Ubuntu 18.04.3 LTS virtual machine. Effective subreads were generated from the raw data using default parameters. Primer removal, demultiplexing, and poly(A) tail/concatemer trimming were performed with *Lima* v1.11.0 with (`-isoseq --peek-guess`) and *isoseq3 refine* (`--require-polya`). Data from the three SMRT cells were then merged. Consensus sequences corresponding to the same transcript were clustered and polished using the *isoseq3 cluster* (`--verbose –use-qvs`). Without a reference transcriptome, HQ isoform transcripts were input into the Cogent v7.0.0 to reconstruct the *de novo* transcriptome following the recommended guidelines. Transcripts were first grouped using minimap2 v2.17 and partitioned into gene families based on k-mer similarity. Each partition was independently reconstructed to generate contigs, building a *de novo P. collini* transcriptome. Transcriptome completeness was evaluated using BUSCO benchmarking against universal single-copy orthologs.

Coding regions within the transcriptome were predicted using TransDecoder v5.5.0. Open reading frames (ORFs) of at least 100 amino acids were considered, with priority given to ORFs with homology to known proteins via BlastP and PFAM searches. Both the standard and alternative genetic codes were applied, and peptide predictions were scored based on PFAM hits. The standard genetic code was selected for the best overall predictions. Functional annotations for the transcriptome were performed using Trinotate v3.2.1, integrating BLASTP v2.3.0+, HMMER/PFAM v3.3.1, SignalP v5, and TMHMM v2 datasets. The resulting *de novo* transcriptome and functional annotation data are available [link].

### Transcription abundance and differential transcript expression analysis

Illumina paired-end reads were aligned to the *de novo P. collini* transcriptome using minimap2 v2.24. Transcript-level abundance and effective transcript lengths were estimated using Salmon v1.13.1 following the recommended guidelines^95^. Salmon detected the library type (`-l A`) and used selective alignment (`--validateMappings`) to identify the potential mapping loci, while correcting fragment-level GC biases (`--gcBias`). Quantifications from individual samples were merged using `salmon quantmerge` to generate a single quantification table, including effective transcript length, Transcript Per Million (TPM), and read counts. The merged quantification table was imported into R via the `tximport` package. Differential transcript expression analysis was computed using `DESeq2`^96,97^, with transcripts pre-filtered to those with at least 10 counts across at least four samples. Each experimental time point was compared to its corresponding non-detentacled negative control group, using the model (design = ∼ time + experiment). Log2 fold changes (LFC>1.5) and significance were computed, with adjusted p-values computed using the Benjamini–Hochberg method to control the false discovery rate (FDR) (adjusted p-value < 0.05). Multiple shrinkage methods (`apeglm` and `ashr`) were applied to refine LFC estimates and only overlapping top hits from both methods were selected. Differentially expressed transcripts were then clustered and analyzed in Python.

### Label-free quantitative mass spectrometry analysis

The *P. collini* cells were concentrated by straining through the 10 μm cell strainer (PluriStrainer) and pelleted by centrifugation at 2000 x RCF for 10 minutes at room temperature. Cell pellets were resuspended in 50 μL of lysis buffer (50 mM Tris-HCl, pH 8, 75 mM NaCl, 8 M guanidine hydrochloride (Gdn-HCl)) and lysed at 95°C for 5 minutes. The lysates were diluted with 50 mM Tris-HCl (pH 8) to reduce the Gdn-HCl concentration below 4 M. Protein concentrations were determined using the Pierce BCA Protein Assay Kit (Thermo Scientific 23227) according to the manufacturer’s instructions. Cell lysates were reduced with 5 mM DTT for 1 hour at 37°C, alkylated with 10 mM iodoacetamide (IAA) in the dark for 1 hour at 25°C, and quenched with 15 mM DTT for 15 minutes. Lysates were further diluted with 50 mM Tris-HCl (pH 8) to reduce the Gdn-HCl concentration below 1 M and digested overnight at 37°C with sequencing-grade modified trypsin (Promega V5113) at a 1:50 protease-to-protein ratio, as determined by the BCA assay. Trypsin-digested samples were desalted using SOLA reversed-phase polymeric cartridges (Thermo Fisher Scientific, 60109-001) before LC-MS/MS analysis.

LC-MS/MS analysis was performed using an UltiMate 3000 RSLCnano system (Thermo Fisher Scientific) coupled to an Exploris 480 mass spectrometer (Thermo Fisher Scientific). 1 µg of sample was loaded onto an Acclaim PepMap C18 100 column (75 µm × 500 mm, 2 µm particle size, 100 Å pore size) with a 120-min gradient. The Exploris 480 was operated with a spray voltage of 2 kV and an ion source temperature of 325°C in positive ion mode. MS1 spectra were collected at 60,000 resolution over a mass range of 350–1,200 m/z with a MS1 AGC target set to 300% with “auto” maximum injection time. MS/MS spectra were acquired at a resolution of 15,000 via HCD fragmentation with a normalized collision energy of 30% for the top 20 peaks with a charge state of 2-6. An isolation width of 1.4 m/z, a dynamic exclusion window of 20 s and a precursor intensity threshold of 5 × 10^3^ was used. The MS/MS AGC target was set to “standard” with a maximum injection time of 22 ms.

Raw LC-MS/MS data files were processed using Proteome Discoverer 2.4 (Thermo Fisher Scientific) with the SEQUEST HT search^98^ . The data was searched against the FASTA database file (RNA-Seq-Pcollini_20210314.fasta). A precursor mass tolerance of 10 ppm and a fragment mass tolerance of 0.02 Da were used. High-confidence peptides were validated using Percolator with a target FDR set to 0.01^99^. Protein identifications of high confidence were determined with an FDR of 0.01 and quantified based on at least one high-confident peptide. Mass modifications in the search included: carbamidomethylation (Cys, +57.021 Da, static), oxidation (Met, +15.995 Da, dynamic), acetylation at protein N-termini (+42.011 Da, dynamic), Met loss at protein N-termini (−131.040 Da, dynamic), Met loss + acetylation at N-termini (−89.030 Da, dynamic). For label-free quantification, peptides were quantified based on precursor abundance intensity, and abundances for each sample were normalized based on total peptide abundance. Proteins were quantified based on the summed normalized intensities of unique and razor peptides, and the summed protein abundances were used to calculate protein abundance ratios comparing starved and fed populations. P-values of abundance ratios were calculated using an ANOVA test based on individual proteins.

### Immunostaining and confocal microscopy

Immunostaining protocols were performed following protocols established for other ciliates, with modifications to better preserve 3D cellular structures more^27,29^. Cells were strain-cleaned and plated in 96-well glass-bottom plates (Cellvis P96-1.5H-N). Cells were fixed directly in the well with a fixative solution (2% (v/v) paraformaldehyde (PFA) and 0.5% (v/v) Triton X-100 in PHEM buffer) at room temperature for 30 minutes. Fixative was then replaced with 0.1 M glycine in PBS and incubated for 30 min at room temperature. Cells were subsequently incubated in the blocking solution (0.2% bovine serum albumin (w/v) in PBS) for 30 min. For centrin staining, cells were first incubated in antibody solution (0.1%(w/v) BSA, 0.1%(v/v) Triton X-100 in PBS) containing a 1:500 dilution of anti-centrin 20H5 mouse monoclonal antibody (Millipore 04-1624) at 4°C overnight. Cells were washed three times for 15 minutes in PBS containing 0.2% (v/v) Triton X-100, then incubated in the antibody solution containing a 1:1000 dilution of Alexa Fluor 568-conjugated anti-mouse IgG antibody (Invitrogen A11004) at room temperature for 1 hour. Cells were washed three times for 15 minutes in PBS with 0.2% (v/v) Triton X-100. For microtubule staining, the cells were incubated in antibody solution containing the Alexa Fluor 488 conjugated anti-alpha-tubulin antibody (Invitrogen 322588) diluted 1:500 at room temperature for 1 hour, followed by three washes of 15 minutes each in PBS with 0.2% (v/v) Triton X-100. Nuclear staining was performed with a 1:1000 dilution of DAPI (Thermo Scientific 62248) for 5 minutes at room temperature, followed by three PBS washes before imaging. For confocal microscopy, the cells were imaged using a Nikon A1R-Si+ confocal microscope and SoRa/W1 spinning disk microscope (Nikon).

### Ultrastructure Expansion Microscopy (U-ExM)

Cells were strain-cleaned and fixed using multiple fixation methods, as *P. collini* exhibited limited cell expansion during U-ExM sample preparation and appeared highly sensitive to fixation methods. For chemical fixation, cells were mixed with an equal volume of 8% paraformaldehyde (PFA) and 2% Triton X-100 (v/v) in PBS, resulting in a final 4% PFA and 1% Triton X-100 (v/v). This fixation method best preserved overall structural integrity, although only partial expansion of *P. collini* cell bodies was achieved. Cells were also fixed with organic solvents such as pre-chilled acetone or methanol at -20°C for 5 minutes. These treatments improved expansion quality, allowing near-complete cell body expansion. However, they caused structural disruption, particularly at the tentacle tips. A mechanical fixation method, high-pressure freezing (HPF), was tested. Concentrated *P. collini* samples (1 μL) suspended in 10% polyethylene glycol (PEG) 3350 were loaded into hexadecane-coated 3 mm carriers and frozen using a Leica EM ICE (Leica Microsystems). Freeze substitution was performed using a Leica automated freeze-substitution system, and the samples were rehydrated according to established protocols before storage in PBS at 4°C for downstream use^3,59,100,101^. Despite improved ultrastructure preservation, HPF did not enhance cell expansion during subsequent U-ExM sample preparation.

U-ExM was carried out following established protocols^3,62,63^, with minor modifications. Fixed cells were first attached to poly-D-lysine–coated coverslips, then cross-linked in an acrylamide/formaldehyde (AA/FA) solution containing 1 and up to 2% acrylamide and 0.7 and up to 1.4% formaldehyde for 12 h at 37 °C. Gelation was performed using a monomer mix composed of 19% (w/w) sodium acrylate (Combi-Blocks, QC-1489), 10% (w/w) acrylamide (Sigma-Aldrich, A4058), and 0.1% (w/w) N,N′-methylenebisacrylamide (Sigma-Aldrich, M1533) in PBS. The reaction was carried out for 1 h at 37 °C in a humid chamber. For denaturation, gels were incubated in denaturation buffer (50 mM Tris, pH 9.0, 200 mM NaCl, 200 mM SDS) for 15 min at room temperature, followed by incubation at 95 °C for 1.5 h. After cooling, gels were expanded by successive exchanges in Milli-Q water until reaching isotropic expansion. The gel diameter was measured to calculate the expansion factor, and all scale bars in images were adjusted accordingly.

For immunostaining of tubulin and centrin, expanded gels were incubated with primary antibodies prepared in PBS containing 3% BSA and either 0.1% Tween-20 or 1% Triton X-100. Incubations were performed overnight at 37 °C. Tubulin was detected using a 1:250 dilution of a rabbit antibody mix against α- and β-tubulin (ABCD_AA345-R and ABCD_AA344-R, ABCD Antibodies SA). Centrin was labelled using a mouse monoclonal antibody (clone 20H5, 04-1624, Sigma-Aldrich) at a 1:300 dilution. After three washes in PBS containing detergent, secondary antibodies were applied at 1:1000 dilution for 4 h at 37 °C. Rabbit primaries were detected with either Alexa Fluor™ Plus 488 (A32790, Thermo Fisher) or Alexa Fluor™ Plus 647 (A32795, Thermo Fisher) donkey anti-rabbit IgG (H+L) antibodies, and the mouse primary was detected with Alexa Fluor™ 568 goat anti-mouse IgG (H+L) (A11004, Thermo Fisher). Gels were washed twice in PBS to remove excess secondary antibodies. For total protein pan-labelling, gels were incubated with Alexa Fluor™ 405 NHS-Ester (A30000, Thermo Fisher) diluted 1:500 in 0.1 M NaHCO₃ buffer (pH 8.3) for 1.5 h at room temperature, followed by two PBS washes and final re-expansion in water. Expanded gels were trimmed to fit, mounted on poly-D-lysine–coated coverslips, and sealed using i-Spacers (IS013, Sunjin Lab). Imaging was performed on a Leica STELLARIS 8 inverted confocal microscope equipped with an APO 40× LWD water immersion objective (1.10 NA, CORR CS2).

### Phenomenological modeling of *P. collini* morphological circuit

We developed a phenomenological model to describe the self-organizing morphological circuit that encodes anisotropic scaling of individual *P. collini* cells into their local optimal tentacle-trap structures. The model represents a cell as a collection of *N* tentacles with lengths {*L*_1_, *L*_2_,…, *L*_N_}, with a finite pool of structure resources *R* (in units of tentacle length) supporting existing tentacle maintenance. Hence, the total resource level of a given cell *R*_tot_ incorporates resources allocated to individual tentacles and the remaining resources not yet incorporated *R*:

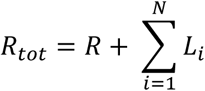

The model integrates deterministic ordinary differential equations (ODEs) for tentacle length dynamics with an event-based probabilistic model for changes in tentacle number (tentacle biogenesis or destruction events), thereby coupling continuous length control and stochastic number jump in a resource-dependent manner.

Inspired by the models of ciliary length control^1,65^ , the growth of each tentacle is positively regulated by the amount of the free resource *R* and negatively regulated by the tentacle’s own length *L*_i_. In the meantime, all tentacles undergo a baseline shrinkage at a constant rate *k*_s_, where the fraction of tentacle length is recycled back to *R*. For each tentacle,

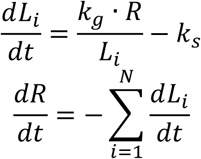

At the steady state, 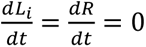, a cell with *N* competing tentacles achieves the same steady-state length *L*_ss_ by symmetry with a free resource level *R*_ss_:

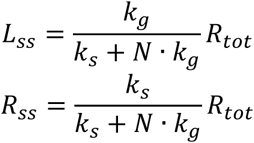

As noticed, the steady-state tentacle length *L*_ss_ depends on the total number of tentacles N sharing the remaining free resource pool *R*_ss_. As N decreases, each tentacle receives a larger share of available resources, yielding a longer steady-state length; vice versa. Because tentacles shorter than ∼15 μm were rarely observed experimentally, we introduce a threshold resource quantity *R*_thre_ required to sustain tentacle formation. We define the optimal tentacle number *N*_opt_as the largest *N* satisfying *R*_ss_ ≥ *R*_thre_. By varying *R*_tot_, we quantified how (*N*_opt_, *L*_ss_) scale with resource availability.

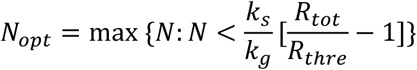

Because the relative magnitude of *R*_thre_to *R*_tot_ determines how readily new tentacles can form, we examined multiple functional relationships between these two parameters, including fixed, linear, reciprocal, power-law, and logarithmic forms. Among all these tested relationships, the logarithmic dependence best reproduced the experimentally observed anisotropic scaling and was used in downstream modeling.

To incorporate stochastic changes in tentacle number, we coupled the continuous resource-dependent length control to discrete probabilistic jumps in tentacle number *N*. Tentacle loss was modeled as a Hill-type stochastic event triggered at each existing tentacle (*L*_i_(*i* = 1, 2, … , *N*)) when the free resource level drops below the threshold ( *R* < *R*_ther_ ). *l* is the Hill coefficient controlling the steepness of the switch, *P*_loss,max_ sets the maximum tentacle destruction probability.

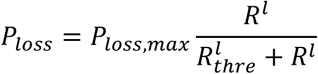

Tentacle formation was implemented as a feed-forward process, where its probability is jointly determined by the free resource pool *R* and a biogenesis factor *Bf*, whose expression is only activated when the free resource level exceeds the threshold (*R* > *R*_thre_ ). *k*_Bf,max_is the maximal synthesis rate of *Bf* and *⍺* defines the saturating response to excess resources. This formulation captures the delayed onset of tentacle biogenesis, starting only when sufficient resources are available.

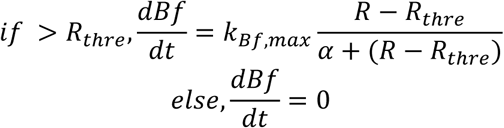

The probability of tentacle formation at a given time step is determined jointly by the *Bf* level and the current free resource pool, each represented by Hill-type switches. *P*_form,max_ represents the maximum tentacle biogenesis probability, *m* and *n* are Hill coefficients, and *K*_Bf_ are the half-saturation constraints.

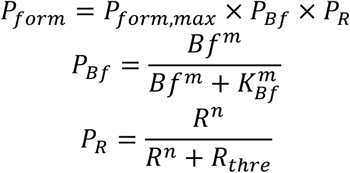

At each time step, probabilities of tentacle biogenesis and destruction are evaluated: tentacle addition occurs with probability *P*_form_ , creating *N*_new_ tentacles following a Poisson distribution of initial length *L*_init_ drawn from the free resource pool *R*; while loss events remove existing tentacles and recycle their length back to *R*. Meanwhile, the lengths of existing tentacles and free resources were dynamically integrated via the deterministic length-control system.

The model enables systematic perturbation of the total resource level *R*_tot_ to study how cells adjust tentacle morphology under different conditions. Feeding was simulated as a stepwise increase in *R*_tot_, and starvation was modeled as a continuous, constant depletion of the resource pool. Synthetic perturbations, such as partial detentacling, can also be simulated as a sudden loss of a subset of tentacles and the associated decrease in the resource.

## Notes

### Competing Interest Statement

The authors have declared no competing interest.

## References.

1. Ishikawa, H., and Marshall, W.F. (2011). Ciliogenesis: building the cell’s antenna. Nat Rev Mol Cell Bio 12, 222–234. 10.1038/nrm3085.

2. Keren, K., Pincus, Z., Allen, G.M., Barnhart, E.L., Marriott, G., Mogilner, A., and Theriot, J.A. (2008). Mechanism of shape determination in motile cells. Nature 453, 475–480. 10.1038/nature06952.

3. Mikus, F., Ramos, A.R., Shah, H., Hellgoth, J., Olivetta, M., Borgers, S., Saint-Donat, C., Araújo, M., Bhickta, C., Cherek, P., et al. (2025). Charting the landscape of cytoskeletal diversity in microbial eukaryotes. Cell. 10.1016/j.cell.2025.09.027.

4. Belyantseva, I.A., Labay, V., Boger, E.T., Griffith, A.J., and Friedman, T.B. (2003). Stereocilia: the long and the short of it. Trends Mol. Med. 9, 458–461. 10.1016/j.molmed.2003.09.008.

5. Chaaban, S., and Brouhard, G.J. (2017). A microtubule bestiary: structural diversity in tubulin polymers. Mol. Biol. Cell 28, 2924–2931. 10.1091/mbc.e16-05-0271.

6. Ridley, A.J. (2011). Life at the Leading Edge. Cell 145, 1012–1022. 10.1016/j.cell.2011.06.010.

7. Rafelski, S.M., and Theriot, J.A. (2024). Establishing a conceptual framework for holistic cell states and state transitions. Cell 187, 2633–2651. 10.1016/j.cell.2024.04.035.

8. Dumont, S., and Prakash, M. (2014). Emergent mechanics of biological structures. Mol. Biol. Cell 25, 3461–3465. 10.1091/mbc.e14-03-0784.

9. Ndlec, F.J., Surrey, T., Maggs, A.C., and Leibler, S. (1997). Self-organization of microtubules and motors. Nature 389, 305–308. 10.1038/38532.

10. McCusker, D. (2020). Cellular self-organization: generating order from the abyss. Mol. Biol. Cell 31, 143–148. 10.1091/mbc.e19-04-0207.

11. Vignaud, T., Blanchoin, L., and Théry, M. (2012). Directed cytoskeleton self-organization. Trends Cell Biol. 22, 671–682. 10.1016/j.tcb.2012.08.012.

12. Yam, P.T., Wilson, C.A., Ji, L., Hebert, B., Barnhart, E.L., Dye, N.A., Wiseman, P.W., Danuser, G., and Theriot, J.A. (2007). Actin–myosin network reorganization breaks symmetry at the cell rear to spontaneously initiate polarized cell motility. J. Cell Biol. 178, 1207–1221. 10.1083/jcb.200706012.

13. Theriot, J.A., and Mitchison, T.J. (1991). Actin microfilament dynamics in locomoting cells. Nature 352, 126–131. 10.1038/352126a0.

14. Gardel, M.L., Schneider, I.C., Aratyn-Schaus, Y., and Waterman, C.M. (2010). Mechanical Integration of Actin and Adhesion Dynamics in Cell Migration. Annu. Rev. Cell Dev. Biol. 26, 315–333. 10.1146/annurev.cellbio.011209.122036.

15. Ponti, A., Machacek, M., Gupton, S.L., Waterman-Storer, C.M., and Danuser, G. (2004). Two Distinct Actin Networks Drive the Protrusion of Migrating Cells. Science 305, 1782–1786. 10.1126/science.1100533.

16. Miller, K.E., and Suter, D.M. (2018). An Integrated Cytoskeletal Model of Neurite Outgrowth. Front. Cell. Neurosci. 12, 447. 10.3389/fncel.2018.00447.

17. Kiddie, G., McLean, D., Ooyen, A.V., and Graham, B. (2005). Biologically plausible models of neurite outgrowth. Prog. Brain Res. 147, 67–80. 10.1016/s0079-6123(04)47006-x.

18. Hjorth, J.J.J., Pelt, J. van, Mansvelder, H.D., and Ooyen, A. van (2014). Competitive Dynamics during Resource-Driven Neurite Outgrowth. PLoS ONE 9, e86741. 10.1371/journal.pone.0086741.

19. Graham, B.P., Lauchlan, K., and Mclean, D.R. (2006). Dynamics of outgrowth in a continuum model of neurite elongation. J. Comput. Neurosci. 20, 43. 10.1007/s10827-006-5330-3.

20. Coyle, S.M. (2020). Ciliate behavior: blueprints for dynamic cell biology and microscale robotics. Mol. Biol. Cell 31, 2415–2420. 10.1091/mbc.e20-04-0275.

21. Jordan, D., Kuehn, S., Katifori, E., and Leibler, S. (2013). Behavioral diversity in microbes and low-dimensional phenotypic spaces. Proc. Natl. Acad. Sci. 110, 14018–14023. 10.1073/pnas.1308282110.

22. Wan, K.Y. (2024). Biophysics of protist behaviour. Curr. Biol. 34, R981–R986. 10.1016/j.cub.2024.07.002.

23. Kysela, D.T., Randich, A.M., Caccamo, P.D., and Brun, Y.V. (2016). Diversity Takes Shape: Understanding the Mechanistic and Adaptive Basis of Bacterial Morphology. PLoS Biol. 14, e1002565. 10.1371/journal.pbio.1002565.

24. Larson, B.T., Garbus, J., Pollack, J.B., and Marshall, W.F. (2022). A unicellular walker controlled by a microtubule-based finite-state machine. Curr. Biol. 32, 3745–3757.e7. 10.1016/j.cub.2022.07.034.

25. Berg, H.C. (1975). Chemotaxis in Bacteria. Annu. Rev. Biophys. Bioeng. 4, 119–136. 10.1146/annurev.bb.04.060175.001003.

26. Leander, B.S. (2020). Predatory protists. Curr. Biol. 30, R510–R516. 10.1016/j.cub.2020.03.052.

27. Coyle, S.M., Flaum, E.M., Li, H., Krishnamurthy, D., and Prakash, M. (2019). Coupled Active Systems Encode an Emergent Hunting Behavior in the Unicellular Predator Lacrymaria olor. Curr Biol 29, 3838–3850.e3. 10.1016/j.cub.2019.09.034.

28. Qin, W., Hu, C., Gu, S., Zhang, J., Jiang, C., Chai, X., Liao, Z., Yang, M., Zhou, F., Kang, D., et al. (2024). Dynamic shape-shifting of the single-celled eukaryotic predator Lacrymaria via unconventional cytoskeletal components. Curr. Biol. 34, 4869–4883.e6. 10.1016/j.cub.2024.09.003.

29. Yanase, R., Nishigami, Y., Ichikawa, M., Yoshihisa, T., and Sonobe, S. (2018). The neck deformation of Lacrymaria olor depending upon cell states. J. Protistol. 51, 1–6. 10.18980/jop.e001.

30. Kourkoulou, A.M., Liu, M., Mathijssen, A.J.T.M., and Juan, G.R.R.-S. (2025). Metachronal wave coordination encodes multimodal swimming in ciliated unicellular predators. bioRxiv, 2025.09.12.675801. 10.1101/2025.09.12.675801.

31. Hara, R., and Asai, H. (1980). Electrophysiological responses of Didinium nasutum to Paramecium capture and mechanical stimulation. Nature 283, 869–870. 10.1038/283869a0.

32. Wessenberg, H., and Antipa, G. (1970). Capture and Ingestion of Paramecium by Didinium nasutum. J. Protozool. 17, 250–270. 10.1111/j.1550-7408.1970.tb02366.x.

33. Bardele, C.F. (1972). A microtubule model for ingestion and transport in the suctorian tentacle. Z. für Zellforsch. Mikrosk. Anat. 126, 116–134. 10.1007/bf00306784.

34. Rudzinska, M.A. (1973). Do Suctoria Really Feed By Suction? BioScience 23, 87–94. 10.2307/1296568.

35. Culbertson, J.R., and Hull, R.W. (1962). Species Identification in Trichophrya (Suctorida) and the Occurrence of Melanin in Some Members of the Genus*. J Eukaryot Microbiol 9, 455–459. 10.1111/j.1550-7408.1962.tb02653.x.

36. Hull, R.W. (1961). Studies on Suctorian Protozoa: The Mechanism of Ingestion of Prey Cytoplasm.*. J. Protozool. 8, 351–359. 10.1111/j.1550-7408.1961.tb01228.x.

37. Hull, R.W. (1961). Studies on Suctorian Protozoa: The Mechanism of Prey Adherence*. J Eukaryot Microbiol 8, 343–350. 10.1111/j.1550-7408.1961.tb01227.x.

38. Kitching, J.A. (1952). Observations on the Mechanism of Feeding in the Suctorian Podophrya. J. Exp. Biol. 29, 255–266. 10.1242/jeb.29.2.255.

39. Terven, J., Córdova-Esparza, D.-M., and Romero-González, J.-A. (2023). A Comprehensive Review of YOLO Architectures in Computer Vision: From YOLOv1 to YOLOv8 and YOLO-NAS. Mach. Learn. Knowl. Extr. 5, 1680–1716. 10.3390/make5040083.

40. Redmon, J., Divvala, S., Girshick, R., and Farhadi, A. (2016). You Only Look Once: Unified, Real-Time Object Detection. 2016 IEEE Conf. Comput. Vis. Pattern Recognit. (CVPR), 779–788. 10.1109/cvpr.2016.91.

41. Budd, G.E. (2021). Morphospace. Curr. Biol. 31, R1181–R1185. 10.1016/j.cub.2021.08.040.

42. Scharf, I., Lubin, Y., and Ovadia, O. (2011). Foraging decisions and behavioural flexibility in trap-building predators: a review. Biol. Rev. 86, 626–639. 10.1111/j.1469-185x.2010.00163.x.

43. James, A., Plank, M.J., and Brown, R. (2008). Optimizing the encounter rate in biological interactions: Ballistic versus Lévy versus Brownian strategies. Phys. Rev. E 78, 051128. 10.1103/physreve.78.051128.

44. Bondoc-Naumovitz, K.G., Laeverenz-Schlogelhofer, H., Poon, R.N., Boggon, A.K., Bentley, S.A., Cortese, D., and Wan, K.Y. (2023). Methods and Measures for Investigating Microscale Motility. Integr. Comp. Biol. 63, 1485–1508. 10.1093/icb/icad075.

45. Dirar, Q., Russell, T., Liu, L., Ahn, S., Dotti, G., Aravamudhan, S., Conforti, L., and Yun, Y. (2020). Activation and degranulation of CAR-T cells using engineered antigen-presenting cell surfaces. PLoS ONE 15, e0238819. 10.1371/journal.pone.0238819.

46. Perry, R.P., and Kelley, D.E. (1970). Inhibition of RNA synthesis by actinomycin D: Characteristic dose-response of different RNA species. J. Cell. Physiol. 76, 127–139. 10.1002/jcp.1040760202.

47. Siegel, M.R., and Sisler, H.D. (1963). Inhibition of Protein Synthesis in vitro by Cycloheximide. Nature 200, 675–676. 10.1038/200675a0.

48. Saxton, R.A., and Sabatini, D.M. (2017). mTOR Signaling in Growth, Metabolism, and Disease. Cell 168, 960–976. 10.1016/j.cell.2017.02.004.

49. Iadevaia, V., Huo, Y., Zhang, Z., Foster, L.J., and Proud, C.G. (2012). Roles of the mammalian target of rapamycin, mTOR, in controlling ribosome biogenesis and protein synthesis. Biochem. Soc. Trans. 40, 168–172. 10.1042/bst20110682.

50. Metzl-Raz, E., Kafri, M., Yaakov, G., Soifer, I., Gurvich, Y., and Barkai, N. (2017). Principles of cellular resource allocation revealed by condition-dependent proteome profiling. eLife 6, e28034. 10.7554/elife.28034.

51. Neilson, K.A., Ali, N.A., Muralidharan, S., Mirzaei, M., Mariani, M., Assadourian, G., Lee, A., Sluyter, S.C. van, and Haynes, P.A. (2011). Less label, more free: Approaches in label-free quantitative mass spectrometry. PROTEOMICS 11, 535–553. 10.1002/pmic.201000553.

52. Sonenberg, N., and Dever, T.E. (2003). Eukaryotic translation initiation factors and regulators. Curr. Opin. Struct. Biol. 13, 56–63. 10.1016/s0959-440x(03)00009-5.

53. Goodson, H.V., and Jonasson, E.M. (2018). Microtubules and Microtubule-Associated Proteins. Cold Spring Harb. Perspect. Biol. 10, a022608. 10.1101/cshperspect.a022608.

54. Chang, C.-C., and Coyle, S.M. (2024). Regulatable assembly of synthetic microtubule architectures using engineered microtubule-associated protein-IDR condensates. J. Biol. Chem. 300, 107544. 10.1016/j.jbc.2024.107544.

55. Mahadevan, L., and Matsudaira, P. (2000). Motility Powered by Supramolecular Springs and Ratchets. Science 288, 95–99. 10.1126/science.288.5463.95.

56. Bombardi, L., Favretto, F., Pedretti, M., Conter, C., Dominici, P., and Astegno, A. (2022). Conformational Plasticity of Centrin 1 from Toxoplasma gondii in Binding to the Centrosomal Protein SFI1. Biomolecules 12, 1115. 10.3390/biom12081115.

57. Lannan, J., Floyd, C., Xu, L.X., Yan, C., Marshall, W.F., Vaikuntanathan, S., Dinner, A.R., Honts, J.E., Bhamla, S., and Elting, M.W. (2024). Fishnet mesh of centrin-Sfi1 drives ultrafast calcium-activated contraction of the giant cell Spirostomum ambiguum. bioRxiv, 2024.11.07.622534. 10.1101/2024.11.07.622534.

58. Chandrasekharan, N.P., Lei, X., Honts, J., Bhamla, S., and Coyle, S.M. (2025). Decoding ultrasensitive self-assembly of the calcium-regulated Tetrahymena cytoskeletal protein Tcb2 using optical actuation. J. Biol. Chem., 110824. 10.1016/j.jbc.2025.110824.

59. Chen, F., Tillberg, P.W., and Boyden, E.S. (2015). Expansion microscopy. Science 347, 543–548. 10.1126/science.1260088.

60. Tillberg, P.W., Chen, F., Piatkevich, K.D., Zhao, Y., Yu, C.-C. (Jay), English, B.P., Gao, L., Martorell, A., Suk, H.-J., Yoshida, F., et al. (2016). Protein-retention expansion microscopy of cells and tissues labeled using standard fluorescent proteins and antibodies. Nat. Biotechnol. 34, 987–992. 10.1038/nbt.3625.

61. Chozinski, T.J., Halpern, A.R., Okawa, H., Kim, H.-J., Tremel, G.J., Wong, R.O.L., and Vaughan, J.C. (2016). Expansion microscopy with conventional antibodies and fluorescent proteins. Nat. Methods 13, 485–488. 10.1038/nmeth.3833.

62. Gambarotto, D., Zwettler, F.U., Guennec, M.L., Schmidt-Cernohorska, M., Fortun, D., Borgers, S., Heine, J., Schloetel, J.-G., Reuss, M., Unser, M., et al. (2019). Imaging cellular ultrastructures using expansion microscopy (U-ExM). Nat. Methods 16, 71–74. 10.1038/s41592-018-0238-1.

63. Mikus, F., Dudin, O., and Dey, G. U-ExM of diverse samples. 10.17504/protocols.io.261gee4qwg47/v1.

64. Ludington, W.B., Ishikawa, H., Serebrenik, Y.V., Ritter, A., Hernandez-Lopez, R.A., Gunzenhauser, J., Kannegaard, E., and Marshall, W.F. (2015). A Systematic Comparison of Mathematical Models for Inherent Measurement of Ciliary Length: How a Cell Can Measure Length and Volume. Biophys. J. 108, 1361–1379. 10.1016/j.bpj.2014.12.051.

65. Marshall, W.F. (2004). CELLULAR LENGTH CONTROL SYSTEMS. Cell Dev. Biol. 20, 677–693. 10.1146/annurev.cellbio.20.012103.094437.

66. Lefebvre, P.A., Asleson, C.M., and Tam, L.-W. (1995). Control of flagellar length in Chlamydomonas. Semin. Dev. Biol. 6, 317–323. 10.1016/s1044-5781(06)80073-3.

67. Marshall, W.F., Qin, H., Brenni, M.R., and Rosenbaum, J.L. (2005). Flagellar Length Control System: Testing a Simple Model Based on Intraflagellar Transport and Turnover. Mol. Biol. Cell 16, 270–278. 10.1091/mbc.e04-07-0586.

68. Marshall, W.F., and Rosenbaum, J.L. (2001). Intraflagellar transport balances continuous turnover of outer doublet microtubules. J. Cell Biol. 155, 405–414. 10.1083/jcb.200106141.

69. Davis, M.H.A. (1984). Piecewise-Deterministic Markov Processes: A General Class of Non-Diffusion Stochastic Models. J. R. Stat. Soc.: Ser. B (Methodol.) 46, 353–376. 10.1111/j.2517-6161.1984.tb01308.x.

70. Mangan, S., and Alon, U. (2003). Structure and function of the feed-forward loop network motif. Proc. Natl. Acad. Sci. 100, 11980–11985. 10.1073/pnas.2133841100.

71. Ma, M., Li, Y., Yuan, Q., Zhao, X., Al-Rasheid, K.A.S., Huang, J., Ma, H., and Chen, X. (2021). New Data Define the Molecular Phylogeny and Taxonomy of Four Freshwater Suctorian Ciliates With Redefinition of Two Families Heliophryidae and Cyclophryidae (Ciliophora, Phyllopharyngea, Suctoria). Front. Microbiol. 12, 768724. 10.3389/fmicb.2021.768724.

72. Hitchen, E.T., and Butler, R.D. (1972). A Redescription of Rhyncheta cyclopum Zenker (Ciliatea, Suctorida)*. J. Protozool. 19, 597–601. 10.1111/j.1550-7408.1972.tb03537.x.

73. Bardele, C.F. (1970). Budding and Metamorphosis in Acineta tuberosa. An Electron Microscopic Study on Morphogenesis in Suctoria. J. Protozool. 17, 51–70. 10.1111/j.1550-7408.1970.tb05158.x.

74. Tucker, J.B. (1974). MICROTUBULE ARMS AND CYTOPLASMIC STREAMING AND MICROTUBULE BENDING AND STRETCHING OF INTERTUBULE LINKS IN THE FEEDING TENTACLE OF THE SUCTORIAN CILIATE TOKOPHRYA. J. Cell Biol. 62, 424–437. 10.1083/jcb.62.2.424.

75. Hickson, S.J., and Wadsworth, J.T. (1909). Dendrosoma Radians, Ehrenberg. J. Cell Sci. *s2-54*, 141–183. 10.1242/jcs.s2-54.214.141.

76. Crocker, A.W., Petrucco, C.A., Guan, K., Wirshing, A.C.E., Ekena, J.L., Lew, D.J., Elston, T.C., and Gladfelter, A.S. (2025). Negative feedback equalizes polarity sites in a multi-budding yeast. Curr. Biol. 35, 3022–3034.e4. 10.1016/j.cub.2025.05.011.

77. Berry, J., Brangwynne, C.P., and Haataja, M. (2018). Physical principles of intracellular organization via active and passive phase transitions. Rep. Prog. Phys. 81, 046601. 10.1088/1361-6633/aaa61e.

78. Alberti, S., and Hyman, A.A. (2021). Biomolecular condensates at the nexus of cellular stress, protein aggregation disease and ageing. Nat. Rev. Mol. Cell Biol. 22, 196–213. 10.1038/s41580-020-00326-6.

79. Li, P., Banjade, S., Cheng, H.-C., Kim, S., Chen, B., Guo, L., Llaguno, M., Hollingsworth, J.V., King, D.S., Banani, S.F., et al. (2012). Phase transitions in the assembly of multivalent signalling proteins. Nature 483, 336–340. 10.1038/nature10879.

80. Linghu, C., An, B., Shpokayte, M., Celiker, O.T., Shmoel, N., Zhang, R., Zhang, C., Park, D., Park, W.M., Ramirez, S., et al. (2023). Recording of cellular physiological histories along optically readable self-assembling protein chains. Nat. Biotechnol. 41, 640–651. 10.1038/s41587-022-01586-7.

81. Lin, D., Li, X., Moult, E., Park, P., Tang, B., Shen, H., Grimm, J.B., Falco, N., Jia, B.Z., Baker, D., et al. (2023). Time-tagged ticker tapes for intracellular recordings. Nat. Biotechnol. 41, 631–639. 10.1038/s41587-022-01524-7.

82. Rajasekaran, R., Chang, C.-C., Weix, E.W.Z., Galateo, T.M., and Coyle, S.M. (2024). A programmable reaction-diffusion system for spatiotemporal cell signaling circuit design. Cell 187, 345–359.e16. 10.1016/j.cell.2023.12.007.

83. Tan, T.H., Liu, J., Miller, P.W., Tekant, M., Dunkel, J., and Fakhri, N. (2020). Topological turbulence in the membrane of a living cell. Nat Phys 16, 657–662. 10.1038/s41567-020-0841-9.

84. Bement, W.M., Leda, M., Moe, A.M., Kita, A.M., Larson, M.E., Golding, A.E., Pfeuti, C., Su, K.-C., Miller, A.L., Goryachev, A.B., et al. (2015). Activator–inhibitor coupling between Rho signalling and actin assembly makes the cell cortex an excitable medium. Nat. Cell Biol. 17, 1471–1483. 10.1038/ncb3251.

85. Chau, A.H., Walter, J.M., Gerardin, J., Tang, C., and Lim, W.A. (2012). Designing Synthetic Regulatory Networks Capable of Self-Organizing Cell Polarization. Cell 151, 320–332. 10.1016/j.cell.2012.08.040.

86. Drubin, D.G., and Nelson, W.J. (1996). Origins of Cell Polarity. Cell 84, 335–344. 10.1016/s0092-8674(00)81278-7.

87. Marshall, W.F. (2020). Pattern Formation and Complexity in Single Cells. Curr. Biol. 30, R544–R552. 10.1016/j.cub.2020.04.011.

88. Lutkenhaus, J. (2012). The ParA/MinD family puts things in their place. Trends Microbiol 20, 411–418. 10.1016/j.tim.2012.05.002.

89. Irazoqui, J.E., and Lew, D.J. (2004). Polarity establishment in yeast. J. Cell Sci. 117, 2169–2171. 10.1242/jcs.00953.

90. Heldt, F.S., Tyson, J.J., Cross, F.R., and Novák, B. (2020). A Single Light-Responsive Sizer Can Control Multiple-Fission Cycles in Chlamydomonas. Curr Biol 30, 634–644.e7. 10.1016/j.cub.2019.12.026.

91. Xu, Z., Chang, C.-C., and Coyle, S. (2024). Synthetic forms most beautiful: engineering insights into self-organization. Physiology. 10.1152/physiol.00064.2024.

92. Milo, R., Shen-Orr, S., Itzkovitz, S., Kashtan, N., Chklovskii, D., and Alon, U. (2002). Network Motifs: Simple Building Blocks of Complex Networks. Science 298, 824–827. 10.1126/science.298.5594.824.

93. Ma, W., Trusina, A., El-Samad, H., Lim, W.A., and Tang, C. (2009). Defining Network Topologies that Can Achieve Biochemical Adaptation. Cell 138, 760–773. 10.1016/j.cell.2009.06.013.

94. Lim, W.A., Lee, C.M., and Tang, C. (2013). Design Principles of Regulatory Networks: Searching for the Molecular Algorithms of the Cell. Mol. Cell 49, 202–212. 10.1016/j.molcel.2012.12.020.

95. Patro, R., Duggal, G., Love, M.I., Irizarry, R.A., and Kingsford, C. (2017). Salmon provides fast and bias-aware quantification of transcript expression. Nat. Methods 14, 417–419. 10.1038/nmeth.4197.

96. Love, M.I., Huber, W., and Anders, S. (2014). Moderated estimation of fold change and dispersion for RNA-seq data with DESeq2. Genome Biol. 15, 550. 10.1186/s13059-014-0550-8.

97. Love, M.I., Anders, S., Kim, V., and Huber, W. (2015). RNA-Seq workflow: gene-level exploratory analysis and differential expression. F1000Research 4, 1070. 10.12688/f1000research.7035.1.

98. Eng, J.K., McCormack, A.L., and Yates, J.R. (1994). An approach to correlate tandem mass spectral data of peptides with amino acid sequences in a protein database. J. Am. Soc. Mass Spectrom. 5, 976–989. 10.1016/1044-0305(94)80016-2.

99. Käll, L., Canterbury, J.D., Weston, J., Noble, W.S., and MacCoss, M.J. (2007). Semi-supervised learning for peptide identification from shotgun proteomics datasets. Nat. Methods 4, 923–925. 10.1038/nmeth1113.

100. Hinterndorfer, K., Laporte, M.H., Mikus, F., Tafur, L., Bourgoint, C., Prouteau, M., Dey, G., Loewith, R., Guichard, P., and Hamel, V. (2022). Ultrastructure expansion microscopy reveals the cellular architecture of budding and fission yeast. J. Cell Sci. 135, jcs260240. 10.1242/jcs.260240.

101. Laporte, M.H., Klena, N., Hamel, V., and Guichard, P. (2022). Visualizing the native cellular organization by coupling cryofixation with expansion microscopy (Cryo-ExM). Nat. Methods 19, 216–222. 10.1038/s41592-021-01356-4.

