## Supplemental Materials for "A self-organizing single-cell morphology circuit optimizes *Podophrya collini* predatory trap structure"

**This document includes:**

Supplemental Figures S1-S5

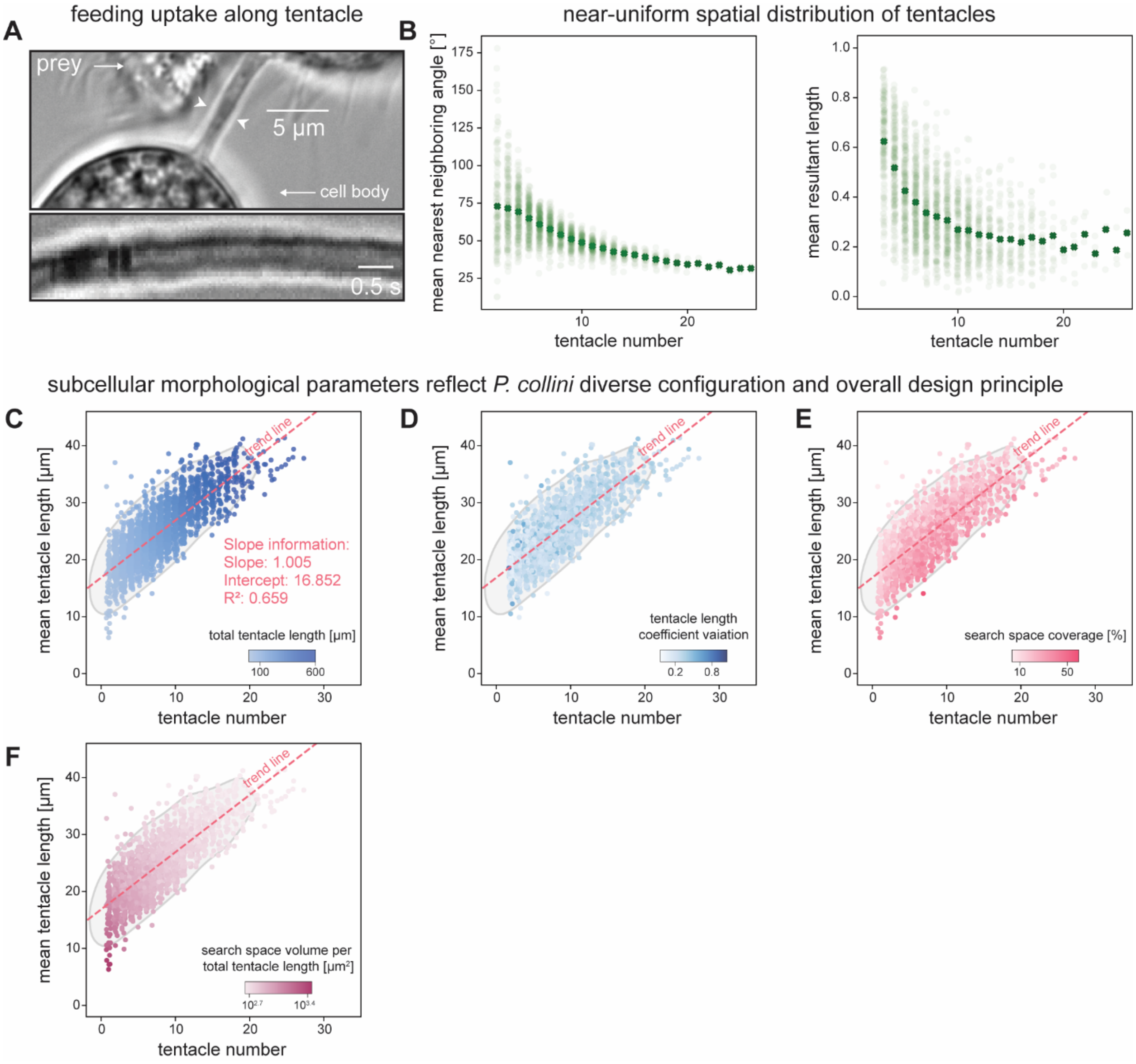


**Figure S1. Subcellular morphological parameters of *Podophrya collini* reflect overarching design principles and diverse structural configurations.**

(A) Representative bright-field DIC images of *P. collini* tentacles during feeding. Top: Prey (*Tetrahymena pyriformis*) cytoplasm is ingested as food vacuoles (white arrows) and transported along the tentacle shaft into *P. collini* cell body during feeding. Bottom: Kymograph showing the movement of a food vacuole along the tentacle shaft over time.

(B) *P. collini* tentacles show a near-uniform spatial distribution. The spatial organization of tentacles in individual cells was quantified using two angular metrics: the mean nearest-neighbor angle (left), representing the average angular spacing between adjacent tentacles, and the mean resultant length (right), which measures the concentration of tentacle directions on the unit sphere. A mean resultant length of 1 indicates perfect alignment, whereas a mean resultant length of 0 indicates uniform distribution.

(C-F): Subcellular morphological parameters illustrate *P. collini's* diverse configurations and underlying geometric principles. Routinely fed populations (n = 2533 cells) were analyzed as the baseline reference. The linear fitting between the tentacle number and mean tentacle length, along with its goodness-of-fit scores, is shown (pink dashed line). Morphological parameters of individual cells were mapped onto the morphospace: (C) integrated tentacle length, (D) coefficient variation of tentacle length, (E) search space coverage relative to the maximized available volume, (F) tentacle cost-efficiency, defined as search volume per unit total tentacle length.

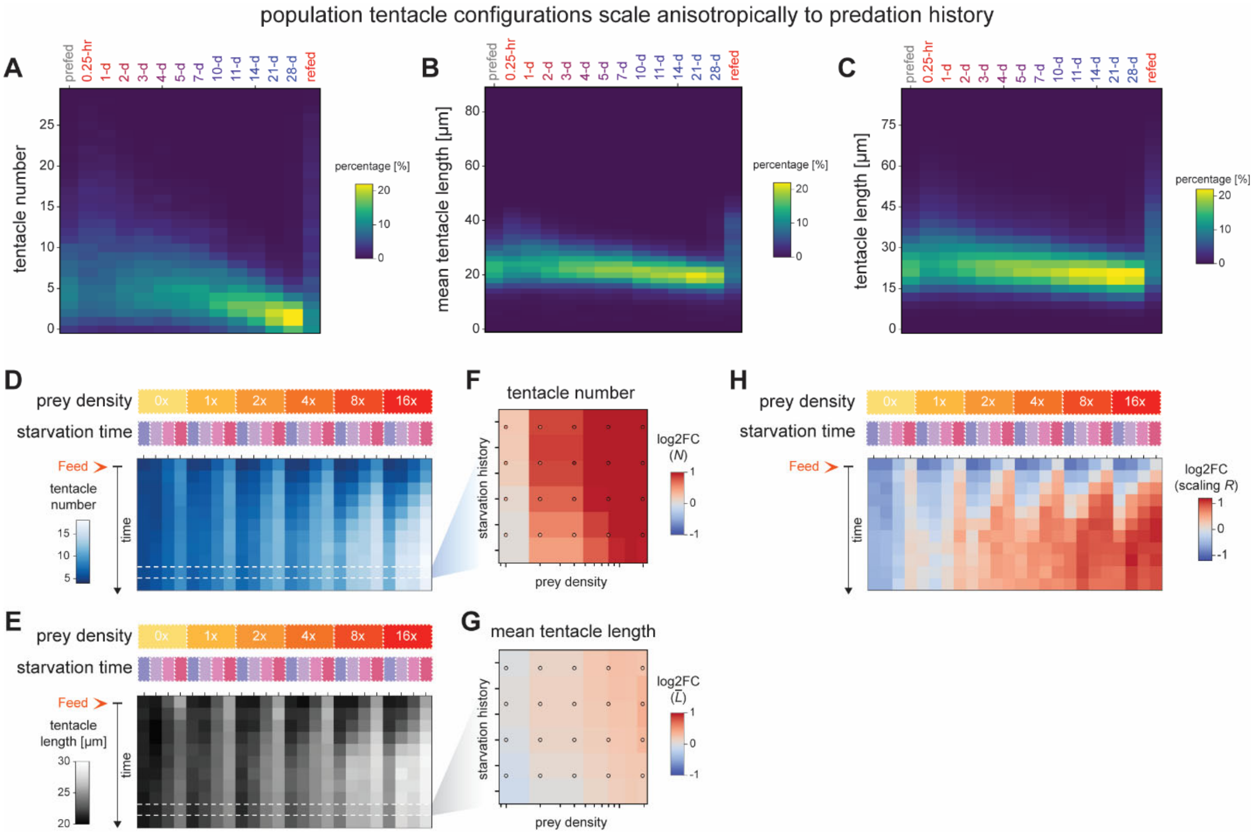


**Figure S2: *P. collini* population tentacle configurations scale anisotropically according to predation history.**

(A-C) Heatmaps showing the distribution of tentacle configuration parameters across the population over time. (A) Tentacle number, (B) mean tentacle length per cell, and (C) individual tentacle lengths.

(D, E) Time-resolved population changes in tentacle configurations across varying feeding contexts ($n_{tot}$ = 243,875 cells). (D) Tentacle number and (E) mean tentacle length.

(F, G) Fitted fold change estimates at 18-hour post-feeding relative to the pre-feeding baseline. (F) Tentacle number and (G) mean tentacle length.

(H) Fold-change of scaling metric *R* over time across different feeding contexts.

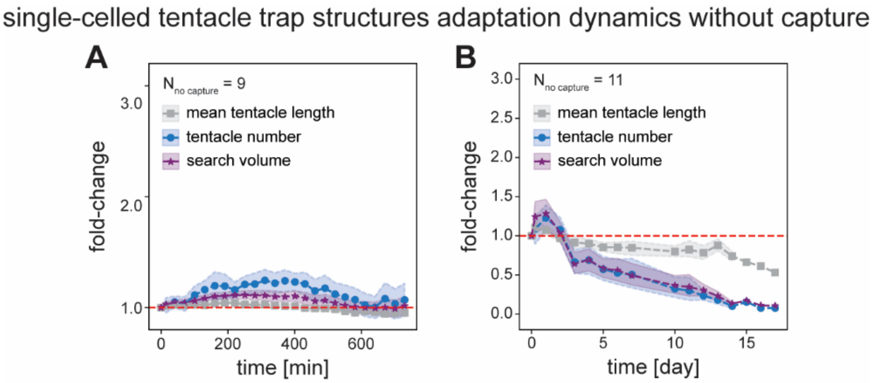


**Figure S3: Individual *P. collini* dynamically tune their tentacle configurations without capture.**

(A, B) Averaged fold-change trajectories of single-cell tentacle configuration parameters over time without prey capture. Symbols represent mean values; shaded regions denote standard deviation. Gray: mean tentacle length; blue: tentacle number; purple: search volume. (A) 12-hour tracking ($n_{nocapture}$ = 9 cells), and (B) 26-day tracking ($n_{nocapture}$ = 11 cells).

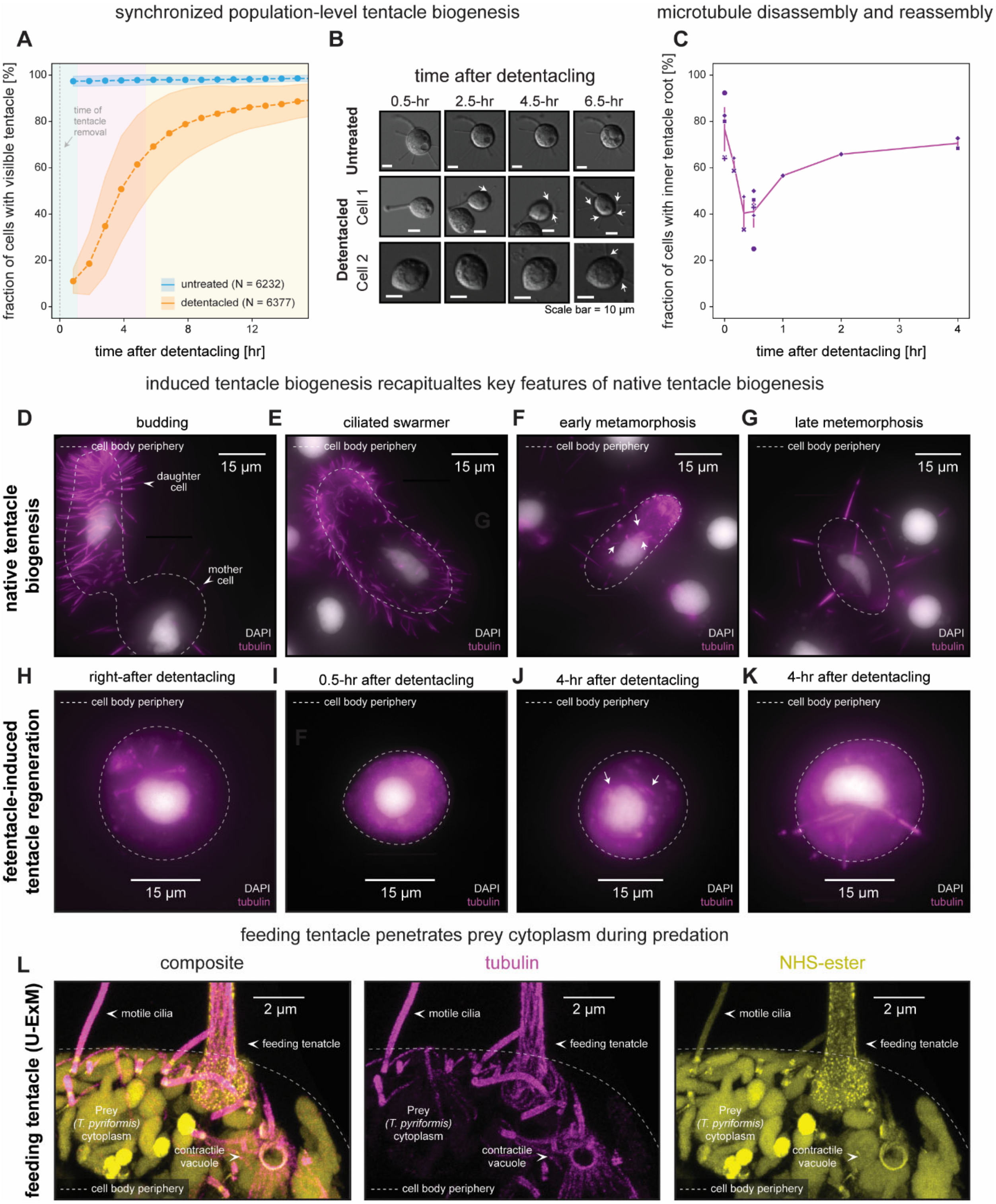


**Figure S4: Tentacle severing synchronously induced *de novo* tentacle biogenesis mirroring native programs.**

(A) Tentacle recovery dynamics over time. Cells were detentacled by mild pressure homogenization and allowed to recover. Tentacle status was scored binarily (with or without tentacles; orange; $n_{treated}$ = 6377 cells; mean ± standard). Unperturbed cells ($n_{untreated}$ = 6232 cells) served as negative controls (blue; mean ± standard). Associated upregulated transcriptional clusters are overlaid with corresponding colors (see Fig. 4J).

(B) Representative bright-field DIC images showing individual cell tentacle status over time in unperturbed and detentacled populations.

(C) Fraction of cells exhibiting inner microtubule roots after detentacling over time. Cells ($N_{tot}$ = 1,060 cells) were immunostained with anti-tubulin and scored under an epifluorescence microscope.

(D-G) Representative epifluorescence images of cells stained with anti-tubulin (magenta) during native tentacle biogenesis at distinct stages: (D) budding, (E) ciliated swarmer, (F) early metamorphosis (transition from ciliated to adult phase), and (G) late metamorphosis. Inner pre-tentacle microtubules are indicated.

(H-K) Representative epifluorescence images of cells stained with anti-tubulin (magenta) during induced tentacle biogenesis following detentacling: (H) immediately after, (E) 30 minutes, and (J, K) 4 hours post-detentacling. Inner pre-tentacle microtubules are indicated.

(L) Representative U-ExM images of a feeding *P. collini* tentacle penetrating prey cytoplasm during feeding. Cells were stained with NHS-ester (yellow) and anti-tubulin (magenta). Prey cilia, contractile vacuoles, and the *P. collini* feeding tentacle are indicated.

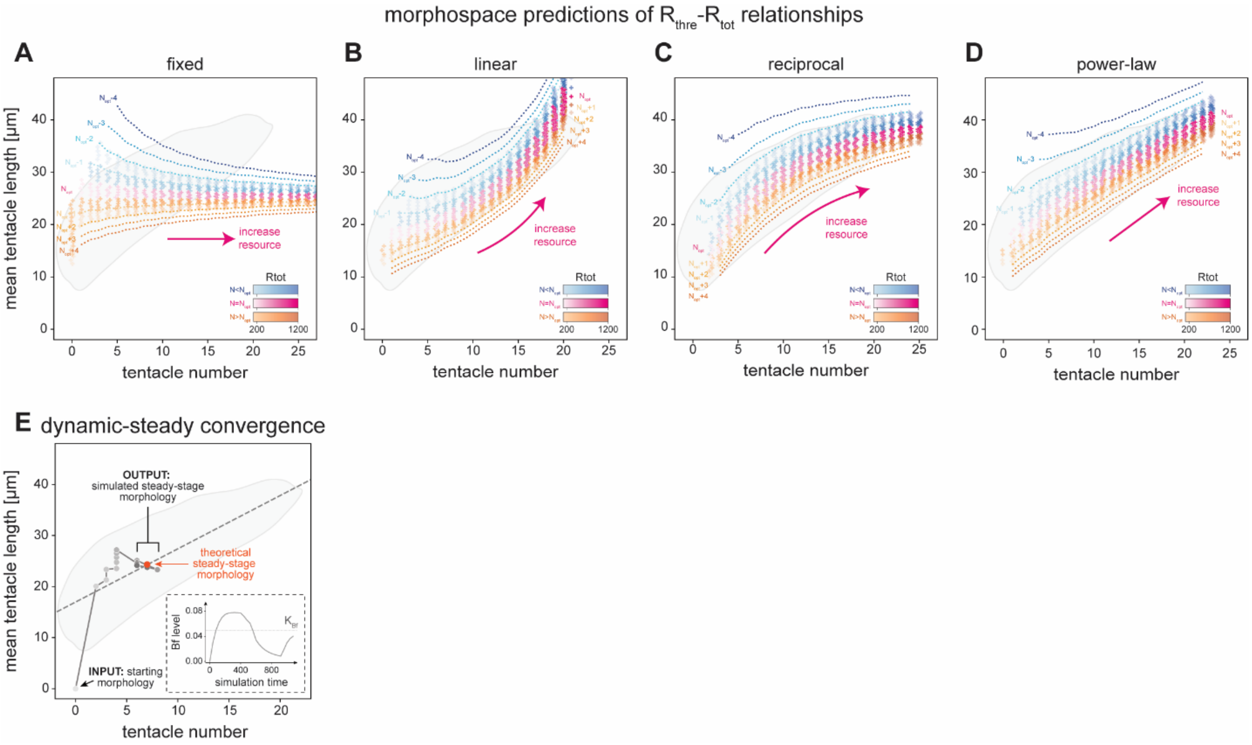


**Figure S5: Mathematical formulation of the *P. collini* cell morphology circuit captures key features of coupled length-number controls.**

(A-D) Predicted tentacle configurations across increasing $R_{tot}$ levels under alternative $R_{thre}$-$R_{tot}$ relationships. The experimentally observed morphospace boundary of routinely fed *P. collini* populations is shown in gray. Steady-state tentacle configurations spanning ($N_{opt}$±4, $L_{ss}$) are mapped onto the morphospace as in Fig. 6B. (A) Constant $R_{thre}$; (B) linear; (C) reciprocal; and (D) power-based $R_{thre}$-$R_{tot}$ relationship.

(E) Tentacle configuration trajectory of a simulated cell reaching its predicted steady state. The cell starts without tentacles at a given $R_{tot}$ and evolves to its predicted end state. The estimated optimal tentacle configuration is indicated in orange. Inset: *Bf* level dynamics over time.
